# Protometabolically generated NADH mediates material properties of aqueous dispersions to coacervate microdroplets

**DOI:** 10.1101/2025.03.13.642830

**Authors:** Rudrarup Bose, Anju Tomar, Daniele Rossetto, Sheref S. Mansy, T-Y Dora Tang

## Abstract

Macromolecular assembly between biomolecules dictates the material state of aqueous dispersions like the cytoplasm. The formation of protein precipitates, fibres, or liquid droplets have been associated with the regulation of biochemical function and disease. However, the effect of small molecules and metabolites on tuning phase behaviour remains underexplored. Here we use the protometabolic reduction of NAD^+^ to NADH by pyruvate to study the effect of NADH production on the phase properties of polyarginine in bicarbonate buffer. Our results show that reduction of NAD^+^ in the presence of polyarginine can tune the material properties of the dispersion between precipitates, homogeneous solution and liquid droplets depending on the buffer concentration. In-situ droplet formation results in 2-3 times higher reaction rate and NADH yield, compared to homogeneous solution. Our results demonstrate how small molecules can significantly impact material properties in aqueous dispersions and the positive effect of heterogeneous dispersions on primitive metabolic reactions.

**Graphical abstract:** 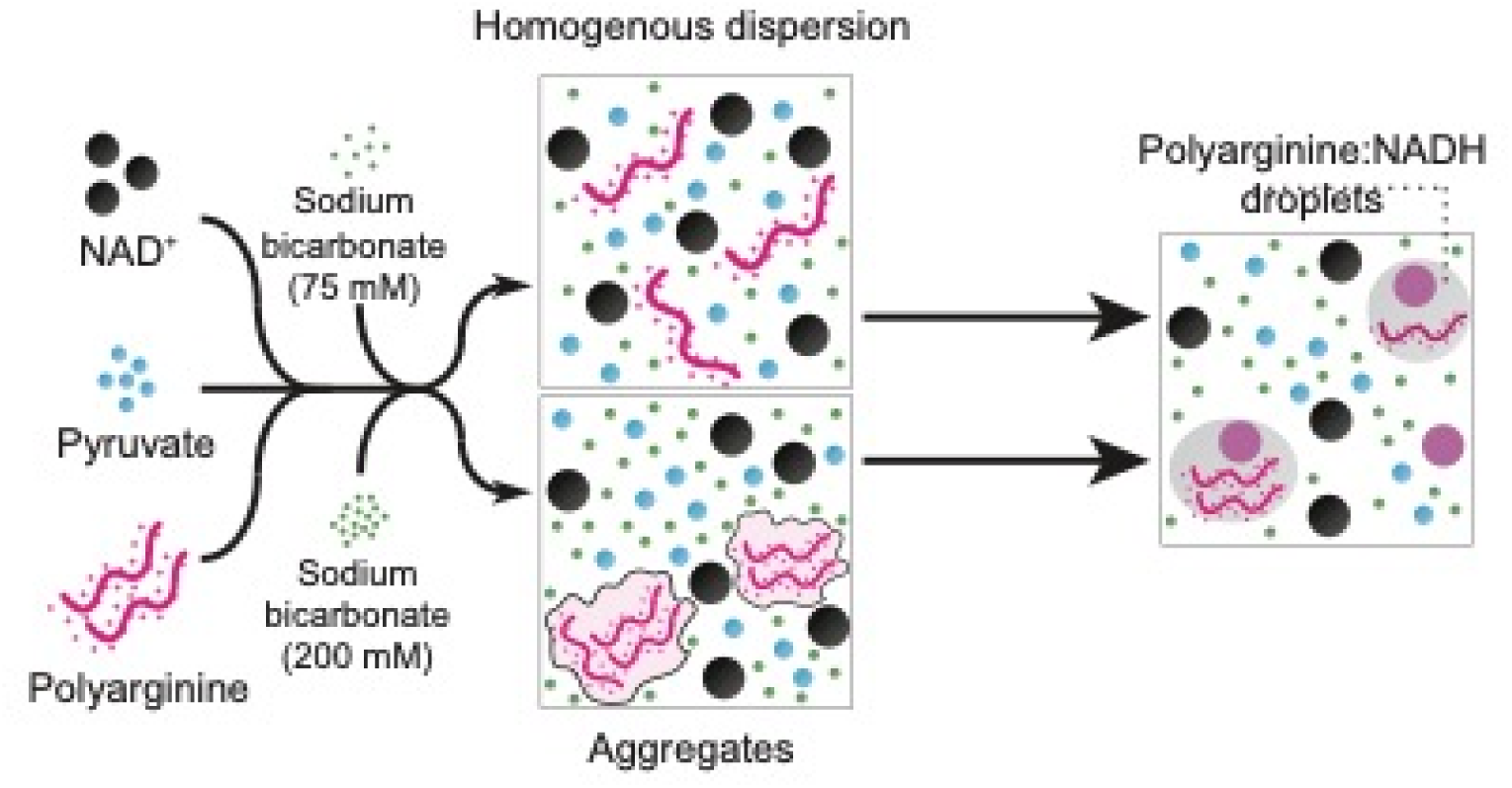

## 1. INTRODUCTION

The biological cell is an active material which exhibits a wide variety of kinetic and thermodynamic material states. These can include, liquid and gel-like droplets or precipitates and fibres where the phase behaviour is dependent on the strength and type of intermolecular interactions; molecular composition and concentration. Further, it has been shown that these material states can be regulated by metabolites^1,2^, underlying biochemical reactions or external stress^3^. Metabolic enzymes have been shown to form a variety of states^4–7^. For example, liver phosphofructokinase (PFK) forms fibres. In yeast cells, glycolytic enzymes and RNA phase separate by liquid-liquid phase separation under stress conditions to regulate glycolysis^8^. In general, reconfiguring metabolic enzymes into compartments or fibres could contain, regulate and protect metabolism^6^. Despite a general understanding that protein droplets can form a variety of material states which can impact biochemical reactions and cellular physiology, there has been little focus on the effect of small molecules or metabolites on the material state of chemically complex aqueous dispersions.

To address this, we sought to use the intrinsic phase behaviour of polyelectrolytes to explore the role of NADH, a key metabolite, for mediating the phases formed by polypeptide complexation. Polyelectrolytes can exhibit complex phase behaviour^9^, forming precipitates or droplets under specific conditions. Polyelectrolyte solutions will precipitate above a critical salt concentration (H-type) or in the case of solutions containing strongly charged polymers solutions, small concentrations of multivalent ions can induce precipitation in a polymer concentration dependent manner (L-type)^10^. Coacervation between two oppositely charged polyelectrolytes provide the thermodynamic driving force for membrane free droplet formation^11,12^ and are considered as relevant models for biological phase separation^13,14^ and questions in the origin of life^15^.

Coacervate dispersions have been shown to exert selective pressures on encapsulated reactions by the partitioning of molecules^16^, by promoting^17,18^, and suppressing reactions^16^ as well as by shifting reaction equilibria^19^ in enzyme catalysed reactions that can lead to shape mediation in peptide-DNA reactions^20^. Non-enzymatic reactions involving small molecules such as the condensation of imine are shown to have higher rates and equilibrium constants in coacervates compared to bulk solution^21,22^. In this case, this was attributed to an upconcentration of reactants in the coacervate droplets. It has recently been shown that the presence of small biological molecules such as ATP which have aromatic and charged functional groups can mediate interactions between proteins to reduce their propensity for aggregation^1,2,23^. To test the ability of metabolites such as NADH to mediate phase behaviour within aqueous dispersions of polyelectrolytes, we used a minimal test-tube system. This comprised the non-enzymatic reduction of NAD^+^ to NADH by the alpha-ketoacid pyruvate^24,25^ in the presence of polyarginine at different buffer conditions (Figure 1a). The redox reaction provides the ability to produce NADH in situ in a non-enzymatic manner that reduces any effect of enzymes on the phase properties of the system that allows direct observation of NADH on phase properties. As polyarginine can precipitate under specific salt conditions (Figure 1b) and forms liquid droplets with NADH^26,27^ the minimal system of bicarbonate buffer, polyarginine and non-enzymatic production of NADH is the ideal system for investigating the effect of NADH on polyelectrolyte phase behaviour. Our results show that in-situ formation of NADH can mediate the material properties of polyarginine from a precipitate phase or homogeneous phase to liquid droplets. Further, the tendency to form liquid droplets will increase the production of NADH compared to homogeneous solutions in the absence of polyelectrolytes.

**Figure 1.**
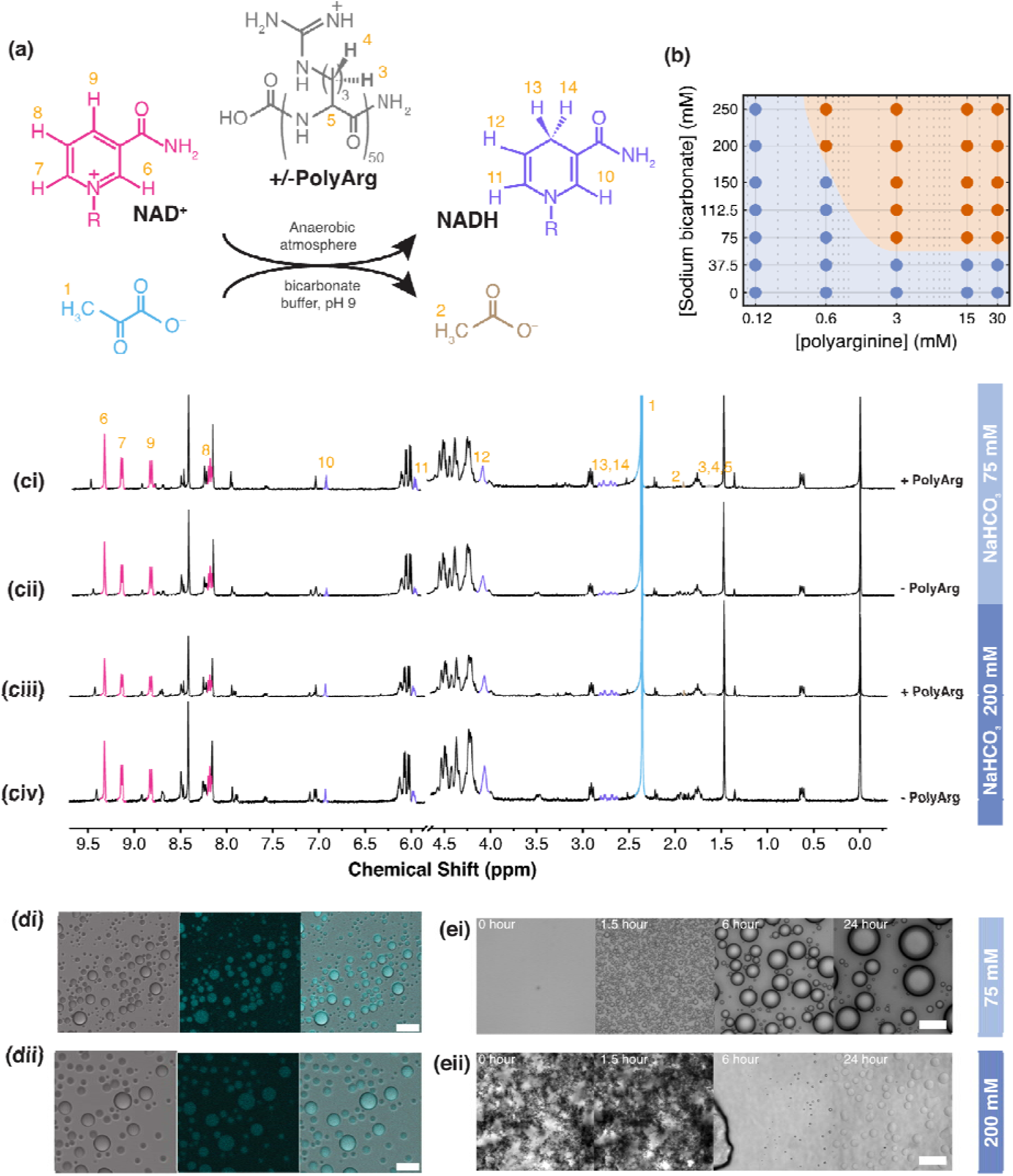
(a) Schematic of reaction, chemical structures and protons identifiable by ^1^H NMR. (b) Phase diagram with semi-log plot of sodium bicarbonate and polyarginine (50 mer concentration (c) ^1^H NMR spectra of reaction mixtures containing 15 mM NAD^+^ 30 mM pyruvate in either 75 mM sodium bicarbonate with (ci) 15 mM of polyarginine or (cii) in the absence of polyarginine or 200 mM sodium bicarbonate (ciii) 15 mM of polyarginine or (civ) in the absence of polyarginine. (d) Fluorescence microscopy imaging of coacervate droplets. bright-field (left), fluorescence (center) and merged (right) images of the reaction with (di) 75 mM and (dii) 200 mM sodium bicarbonate buffer, showing droplets formed after 6 hours from the start of the reaction (scale bar = 20 µm). The fluorescence images were acquired by illuminating with a light source of 355 nm and detecting emission between 449 nm and 506 nm. (e) time-lapse bright field images of the reaction with polyarginine at (ei) 75 mM and (eii) 200 mM sodium bicarbonate buffer show a transition to a droplet state.

## 2. EXPERIMENTAL SECTION

### 2.1. Materials

β-nicotinamide adenine dinucleotide disodium salt (NAD^+^), β-nicotinamide adenine dinucleotide, reduced disodium salt (NADH), L-arginine monohydrochloride (arginine), L-lysine (lysine), NAD^+^/NADH quantitation kit, sodium pyruvate, and toluene were purchased from Sigma Aldrich USA. Hydrochloric acid (HCl) and sodium hydroxide (NaOH) was purchased from Merck Millipore USA. Poly-L-arginine hydrochloride (polyarginine) and poly-L-lysine hydrochloride (polylysine) was purchased from Alamanda Polymers USA. 2-[methoxy(polyethyleneoxy)propyltrimethoxysilane, 6-9 PEG units (PEG-silane) from ABCR, guanidine thiocyanate (GuSCN) from PanReac AppliChem Germany, sodium trimethylsilylpropanesulfonate (DSS) from Tokyo Chemical Industry Japan, sodium bicarbonate (NaHCO_3_) from Merck Germany, Hellmanex III from Hellma and deuterated water (D_2_O) from Deutero GMBH Germany was purchased.

### 2.2. Preparation of reaction mixtures

In all experiments, reaction mixtures were prepared to a final concentration of 15 mM NAD^+^ and 30 mM sodium pyruvate with either 75 mM or 200 mM NaHCO3 buffer at pH 9 with and without 15 mM polyarginine. In addition, experiments were undertaken with 15 mM NAD^+^, 30 mM sodium pyruvate, 75 mM NaHCO_3_ with either 15 mM or 150 mM of monomer arginine.

MiliQ water and 500 mM NaHCO_3_ at pH 9.0 (adjusted with 7.5 M sodium hydroxide) were degassed using standard protocols. Solutions were placed in a Schlenk line and frozen on a mixture of acetone and dry-ice. Oxygen was removed by a vacuum pump followed by thawing on warm water. To ensure that the maximum amount of oxygen had been removed, the procedure of freezing, vacuum and thawing was repeated 5 times. The solutions were then stored under nitrogen flux for up to a week. At the time of the experiments, solutions were transferred from the Schlenk line into the glove box in a Teflon capped vial, preconditioned with nitrogen flux.

Stock solutions (A-D) (Table 1) were prepared in a glove box with a nitrogen flow and oxygen levels below 0.1% (measured using Okolab LEO hand-held meter) to ensure anaerobic conditions. NAD^+^, sodium pyruvate, polyarginine and arginine solids were weighed into micro-centrifuge tubes and transferred into the glove chamber along with degassed Milli-Q water and 500 mM sodium bicarbonate buffer (pH 9.0).

**Table 1.** List of stock solutions for reaction mixture.

| Solution | Chemical mixture | Aqueous solution |
| --- | --- | --- |
| A | 25 mM polyarginine or monomer arginine | Mili-Q water |
| B | 250 mM polyarginine or monomer arginine | Mili-Q water |
| C | 75 mM pyruvate, 37.5 mM NAD<sup>+</sup> | 187.5 mM sodium bicarbonate buffer (pH 9) |
| D | 75 mM pyruvate, 37.5 mM NAD<sup>+</sup> | 500 mM sodium bicarbonate buffer (pH 9) |

To prepare 100 µL of reaction mixture with 75 mM NaHCO_3_ buffer, 40 µL of solution C was added to 60 µL of solution A / solution B / or degassed water for a reaction mixture containing 15 mM (solution A) or 150 mM (solution B) of polyarginine or arginine, or a reaction mixture free from peptides. To prepare reaction mixtures containing 200 mM NaHCO_3_, 40 µL of solution D was added to 60 µL of solution, A, B, or degassed water to give final concentration of 15 mM (solution A) or 150 mM (solution B) of poly peptide or amino acid or a reaction mixture free from polypeptide or amino acids.

### 2.3. Characterisation of reaction by NMR

Samples were prepared as described previously under anaerobic conditions. To produce the 600 µL of sample required for NMR experiments the volumes were scaled up. For experiments undertaken in the absence of (poly) amino acids, 500 mM NaHCO_3_ buffer was prepared with 10% D_2_O water and 1 mM of sodium trimethylsilylpropanesulfonate (DSS) and degassed as previously described. For reactions that contained (poly) arginine, the reaction mixture was prepared as previously described. At specified time points the sample was diluted with 10 mM DSS prepared in D_2_O to prepare a final reaction sample containing 10% of D_2_O and 1 mM DSS.

Samples were loaded into a NMR TUBE (Boroeco-5-7, Deutero) and inserted into a BRUKER NanoBay 400 MHz NMR spectrometer. ^1^H NMR spectra were obtained with water suppression using the Bruker’s noesygppr1d pulse program with 16 scans (fid size: 32768). In summary, the following settings were used: spectral width: 15.9794 ppm / 6393.862 Hz, acquisition time: 2.5624576 s, fid resolution: 0.390250 Hz, filter width: 4032 kHz, transmitter frequency offset: 4.7 ppm / 1880.61 Hz, receiver gain: 32, dwell time: 78.2 µs, DSP firmware filter: rectangle, digitization mode: baseopt, digitizer resolution: 22, homodecoupling duty cycle: 20%, oversampling during homodecoupling: 1, maximum variation of a delay: 5%, first step for PL switching: -6 dB, step width for PL switching: 0.1, MAS rotation: 4200 Hz, Probe temperature: 298 K, Temperature on channel: 300 K, gradient temperature: 300 K, number of wobble steps: 1024). The NMR spectra were analyzed using Mestrelab Mnova software. The spectrum was adjusted using DSS as the reference, δ 0.00 (s, 9H). The ^1^H NMR spectra was compared against the assignable hydrogens in the ^1^H NMR spectrum of pure NADH, NAD^+^, pyruvate, acetate and polyarginine, i.e., NADH, (Adenosine-P-P-Nicotinamide-H): ^1^H NMR (400 MHz, H2O with 10% D2O) δ 2.72 (dd, *J* = 14.3, 7.9 Hz, 2H), 5.97 (d, *J* = 8.2 Hz, 1H), 6.93 (s, 1H). NAD^+^, (Adenosine-P-P-Nicotinamide^+^): ^1^H NMR (400 MHz, H2O with 10% D2O) δ 8.18 (t, *J* = 7.2, 7.2 Hz, 1H), 8.82 (d, *J* = 8.1 Hz, 1H), 9.13 (d, *J* = 6.3 Hz, 1H), 9.32 (s, 1H). Pyruvate, (O=C-CH_3_)-C(O^-^): ^1^H NMR (400 MHz, H2O with 10% D2O) δ 1.47 & 2.36 (s, 3H). Acetate, (O=C)-(CH_3_): ^1^H NMR (400 MHz, H2O with 10% D2O) δ 1.91 (s, 3H). Polyarginine, (-NH-CH(CNH(NH_2_))-(CH_2_)_5_-NH-C(O)-)_n_: ^1^H NMR (400 MHz, D2O) δ 1.97 – 1.55 (br).

### 2.4. Imaging of in-situ droplet formation

Time-resolved widefield and confocal brightfield and fluorescence microscopy was used to image the reaction mixture. To ensure strict anaerobic conditions, 100 µL of sample was loaded into 18-well chamber slide (Ibidi, Germany) in a glove box (Bel-Art Techni-Dome 360° Glove Chamber) that was adhered to a pegylated cover slip. The 18-well slide was then sealed with an adhesive clear microplate sealing sheet (Thermo Scientific, USA). The slide was further sealed with high vacuum grease (Dow, USA) (Figure S1). The microscope slide was loaded onto Pecon Incubator HF 2000 and imaged at 1% oxygen level, controlled through Okolab CO2-O2 Unit-BL [0-10,1-18]. Images were taken every 15 mins, using Zeiss Axiovert 200M wide-field microscope equipped with Andor Zyla PLUS monochrome sCMOS camera with 6.5 μm dexel size and 20x/0.4 LD A-Plan, Air, Ph2, Zeiss objective (product ID: 421051-9910-000). Multi-well imaging was conducted on 6 wells with 15 ms exposure, gain = 3 and digitizer = 200 MHz. Z-stacks were obtained with step size of 3 µm over a range of 80 µm.

To image NADH^28^ inside the droplets, fluorescence microscopy was conducted using Zeiss LSM 880 airy inverted confocal microscope with Plan-Apochromat 20x/0.8 M27. For illumination of NADH, a 355 nm laser source, with laser power 1.2 µW (0.375 µW/cm^2^), master gain 700 and digital gain 1, was used in conjunction with MBS355 beam splitter. The detection filter was set between 449 nm and 506 nm. Bright field images were obtained using transmission detection module (T-PMT) when the sample was illuminated with a 561 nm laser source, with laser power 6.7 µW (2.075 µW/cm^2^), master gain 250, digital gain 1, in conjunction with MBS488/561 beam splitter. For imaging 30.5 µm pinhole size along with 7 µs scan time, averaging of 4 and bit depth of 8 was used.

### 2.5. Phase diagram characterisation

To determine the phase behaviour of NAD^+^ and NADH with polyamino acids we characterized the phase diagrams of polyarginine (0.15, 1.5, 7.5, 15 and 30 mM) with NAD^+^ or NADH (0.15, 1.5, 7.5, 15 and 30 mM) in either 75 mM or 200 mM NaHCO_3_ buffer. To do this, the following stock solutions were prepared: 60 mM NAD^+^ in 150 mM NaHCO_3_ buffer (pH 9); 60 mM NAD^+^ in 400 mM NaHCO_3_ buffer (pH 9); 60 mM NADH in 150 mM NaHCO_3_ buffer (pH 9); 60 mM NADH in 400 mM NaHCO_3_ buffer (pH 9) and 60 mM polyarginine in MiliQ (∼pH 7). In all instances the pH of NaHCO_3_ was adjusted to pH 9 using NaOH unless otherwise stated.

In addition, the phase behaviour of arginine (15 mM) was determined with NADH (0.5, 1, 5 mM) in 75 mM and 200 mM NaHCO_3_. To do this stock solutions of 30 mM arginine were prepared in sodium bicarbonate buffer (75 mM and 200 mM, pH 9). All stock solutions were stored at -20 °C until further use. The monomer concentration of the polypeptide was reported as the concentration of the polypeptide solution.

To prepare samples for phase behaviour characterization, stock solutions were diluted in their respective buffers or MiliQ water to enable a 1:1 volume mixing between polyarginine or arginine with NAD^+^ or NADH to achieve the final concentration. For example, to prepare samples containing 0.15 mM polyarginine, 0.15 mM NADH and 75 mM NaHCO_3_: 60 mM polyarginine in MiliQ water was diluted to 0.30 mM polyarginine with MiliQ water. Solutions containing 60 mM NADH in 150 mM NaHCO_3_ were diluted to 0.30 mM NADH with 150 mM NaHCO_3_. The solution of polypeptide or peptide was mixed with NADH or NAD^+^ at equal volumes to produce 20 µL of polyarginine / NADH / NaHCO_3_ solutions.

Widefield microscopy imaging was used to determine the phase state of polypeptide with NAD^+^/NADH in either 75 mM or 200 mM NaHCO_3_. To do this, 10 µL of sample was loaded into a capillary slide formed by strips of parafilm, sandwiched between a glass slide and a pegylated coverslip^29^. Cover slips were pegylated to prevent droplet wetting on the surface. In brief, coverslips were first cleaned with Hellmanex III (Hellma) and then with distilled water by sonicating for 5 minutes in each solution. The wash steps were repeated 3 times and the cover-slips were then fully submerged into a well-mixed solution of 100 ml toluene, 460 mg of PEG silane and 160 μL of 36% hydrochloric acid. The cover-slips were left for 18 hours with constant stirring and covered to prevent evaporation. The cover-slips were then rinsed with pure toluene, then two times with pure ethanol and distilled water respectively. Finally, cover-slips were dried with nitrogen and stored under vacuum until use.

Once the sample was loaded into the capillary slide, the samples were imaged with a Zeiss Axiovert 200M wide-field microscope equipped with Andor Zyla 4.2 (VSC-02370) camera with 6.5 μm dexel size and a Zeiss 100x/1.3 oil Plan-Neofluar Ph3 M27 objective. To determine the phase of the sample, images were analyzed by eye to determine whether droplets or precipitates were present. To image samples that did not contain precipitates or droplets, 0.2 µL of 0.02% solution of micrometer sized carboxylate beads (Polybead, Polysciences Europe GmbH, product ID: 08226-15), prepared in MilliQ water, was added to 10 µL of mixture and imaged using bright field microscopy to locate the surface of the cover slip.

### 2.6. Determination of fraction of NADH produced

To quantify NADH formation as a function of time, we used a commercially available dual enzyme assay (NAD^+^/NADH Quantitation kit, Sigma Aldrich, USA) which measures the relative amount of NADH compared to NAD^+^ and NADH (Figure S2).

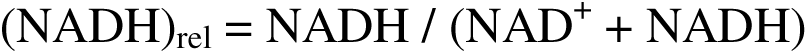

To obtain the relative amount of NADH in the reaction mixture, the reaction mixture was prepared as previously described and incubated under anaerobic conditions. At the time of measurement 4 M guanidine thiocyanate (GuSCN) was added to the reaction mixtures at 1:1 volume to dissolve the coacervate droplets at a final concentration of 2 M GuSCN.

To obtain the amount of NADH in the solution, half of the reaction mixture containing 2 M GuSCN was diluted 1000-fold in NAD^+^ extraction buffer (supplied in the NAD^+^/NADH quantitation kit) and incubated for 30 minutes at 60°C to degrade all NAD^+^. The NAD^+^/NADH quantification assay was then performed according to the manufacturer’s instructions.

To obtain the total amount of NAD^+^ and NADH, the other half of the reaction mixture containing 2 M GuSCN was diluted 10,000 fold in the NAD^+^ extraction buffer and then assayed with the NAD^+^/NADH quantification kit according to the manufacturer’s instructions.

For both reaction mixtures, 25 µL of the diluted reaction mixture was loaded into a 96-well microplate (F-bottom, clear, Greiner Bio-One, Austria) along with 50 µL of enzyme mix. After 5 minutes of incubation, 5 µL of substrate II was added to achieve a total volume of 80 µL. After the colour had developed, 5 µL of stop solution was added as described by the manufacturer’s instructions. With each experiment, a control experiment was performed with known concentrations of NADH (0, 0.4, 0.8, 1.2, 1.6, 2 µM) as provided by the manufacturer. 25 µL of the NADH standards were incubated with 50 µL of enzyme mixture in a well plate reader alongside the reaction samples. Once the absorbance had reached a value between 1.0 to 1.5, the reaction mixture was quenched with the 5 µL of stop solution from the assay kit and the samples were measured by absorbance using a TECAN Spark 20M well plate reader at room temperature (Figure S3a).

The calibration curve was used to determine the absolute quantity of NADH and (NAD^+^ + NADH) from each of the separated reaction mixtures. The concentrations of (NADH) and (NAD^+^ + NADH) were scaled relative to the dilution factors. 3 repeats were undertaken for each time point and each sample.

(NADH)_rel_ was plot as a function of time and the values of (NADH)_rel_ for the first 8 hours was fit to a straight-line without intercept, using MATLAB, to obtain the initial rate of the reaction (Figure S3b). The standard deviation was reported as the error associated with the slope. This was calculated by dividing the range of the 95% confidence interval by 3.92 (since the 95% confidence interval of a prediction is equal to the predicted value ± 1.96 times the standard deviation).

## 3. RESULTS AND DISCUSSION

We first tested the electron transfer reaction between NAD^+^ (15 mM) and pyruvate (30 mM) in sodium bicarbonate buffer 75 mM and 200 mM, at pH 9 as previously described^24^, in the presence of polyarginine (15 mM) and at two different buffer conditions. After 24 hours, all reactions were centrifuged, the supernatant was removed and loaded into an NMR tube with 10% D_2_O for analysis by NMR spectroscopy (Figure 1, Figure S4-S11). Comparison of the reaction mixtures, in the absence and presence of polyarginine (50 mer) and at 75 mM and 200 mM Sodium bicarbonate buffer confirmed the production of NADH and acetate with polyarginine and at 75 mM (low) and 200 mM (high) carbonate concentrations (Figure 1c). To determine the phase state of the reaction mixture with polyarginine, we used optical microscopy to image the reaction after 6 hours. Confocal microscopy images showed the presence of microdroplets that were fluorescent upon UV excitation λ_exc_ = 340 ± 30 nm and λ_emi_ at 450 ± 50 nm. We observed fluorescence emission commensurate with NADH within the microdroplet (Figure 1d). In this case NADH could form the structural component (scaffold) of the coacervate or act as the client by partitioning into the droplet. Given that polyarginine will precipitate above 75 mM of sodium bicarbonate (Figure 1b), we used time-resolved widefield optical microscopy to detect any phase change in the dispersion during the course of the reaction. At 200 mM sodium bicarbonate, the reaction mixture in the presence of polyarginine showed precipitates which transitioned to droplets within 24 hours (Figure 1e). In comparison, at lower sodium bicarbonate concentration (75 mM) there was a homogeneous dispersion at the start of the reaction which then transitioned to droplets within 2 hours of the reaction (Figure 1e). Taken together, the results confirm that non-enzymatic electron transfer reactions can take place in the presence of polyarginine and that the generation of products could be responsible for a change in phase properties within the reaction mixture that could be attributed to NADH production.

To confirm that production of NADH was responsible for the observed phase transitions, we mapped the thermodynamic phase diagrams of polyarginine (0-30 mM) with NADH (0-30 mM) in 75 mM and 200 mM sodium bicarbonate (Figure 2 and S14-S16). The results confirm that NADH can form droplets with polyarginine in both of these conditions. Interestingly, the phase diagrams show a precipitate region towards the bottom right of the phase diagram (low NADH concentration and high polyarginine concentration) (Figures 2a). This phase region is larger at 200 mM sodium bicarbonate compared to 75 mM bicarbonate. The phase diagrams show that increasing the NADH concentration will lead to a transition between a precipitate phase and a droplet phase. It is interesting to note that whilst the phase diagrams show a precipitate at 75 mM bicarbonate and 15 mM polyarginine, no precipitates were observed in the reaction mixture that contained pyruvate immediately after mixing indicating that pyruvate will affect the phase behaviour of the overall system (Figure 1ei).

**Figure 2.**
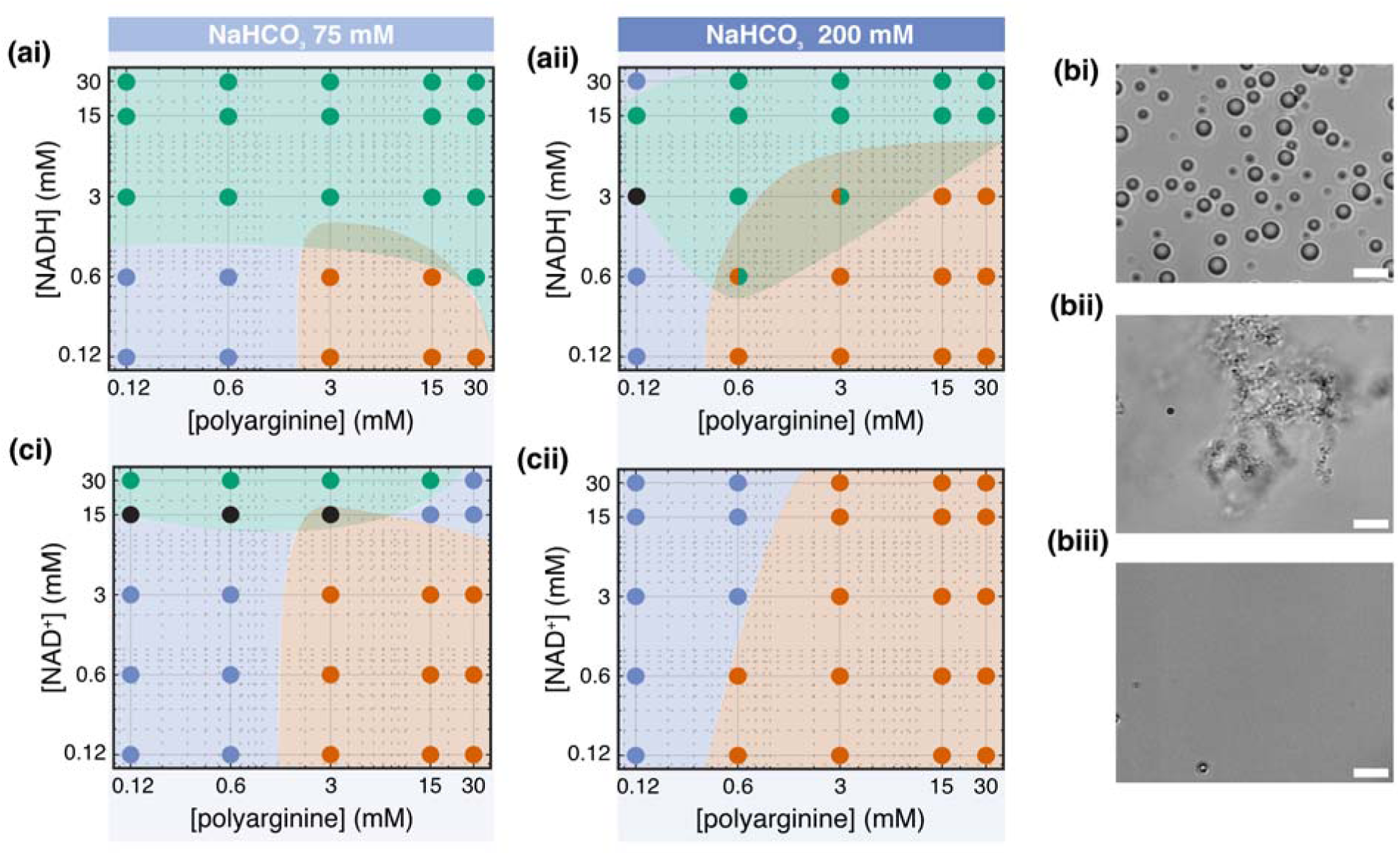
Phase behaviour of polyarginine and NAD^+^/NADH (a) shows the phase diagram of 50-mer polyarginine with NADH in 75 mM (ai) and 200 mM (aii) sodium bicarbonate, respectively, in a log-log plot. (b) Representative images of droplets (15 mM polyarginine, 3 mM NADH in 75 mM sodium bicarbonate) (i), precipitate (15 mM polyarginine, 15 mM NAD^+^ in 200 mM sodium bicarbonate) (ii) and a homogeneous phase (15 mM polyarginine, 15 mM NAD^+^ in 75 mM sodium bicarbonate) (iii). Scale bar = 5 µm. (c) phase behaviour of 50-mer polyarginine with NAD^+^ in 75 mM and 200 mM sodium bicarbonate, respectively, in a log-log plot. Blue, red and green markers represent dissolved solutions, precipitates and droplets. Conditions for which we were unable to determine whether droplet or precipitate were present with certainty are marked black.

Further, the phase diagrams show that mixtures of polyarginine, NADH and buffer can lead to three phase regions (precipitates, droplets and homogeneous phases (Figure 2b)). In order to decipher the stabilizing factors for this diverse phase behaviour we further mapped the phas diagrams of polyarginine in sodium bicarbonate buffer (75 and 200 mM) with NAD^+^ (Figure 2c). At 200 mM of sodium bicarbonate, we observe two phase regions (a homogeneous phase and a precipitate phase), at lower buffer conditions (75 mM) we observe three phases: anhomogeneous phase; a precipitate phase; and a droplet phase. The droplet phase is stabilized at high NAD^+^ concentrations. Comparing phase diagrams with NAD^+^ and NADH shows that NADH increases the droplet phase region even at high buffer conditions.

In summary, the phase diagrams show that increased buffer concentration leads to stabilization of precipitates that could be driven by hydrophobic interactions^30^. On the other hand, NAD^+^/NADH with polyarginine favours droplet formation with NADH having a stronger effect than NAD^+^. As NADH is the reduced form of NAD^+^, NADH is more negatively charged, suggesting that increased electrostatic interactions stabilize the polyarginine droplets. Taken together, this could suggest that in multi-component systems different interactions stabilize different phases. In this instance, hydrophobic interactions at high salt and polypeptide concentrations drive precipitate formation whilst electrostatic interactions lead to droplet formation. The balance in the interactions based on concentration could mediate material states within aqueous dispersions.

Given that we observe different materials properties we next sought to determine the effect of the material states on the kinetics of NADH production. To do this, we measured the fraction of NADH (of the total NAD^+^ and NADH) as a function of time. This was done using a commercially available biochemical assay that was undertaken on the NAD^+^/NADH reaction (Figure S2-3, S17) at 75 mM and 200 mM sodium bicarbonate in the absence and presence of polyarginine (15 mM). Our results show that after 24 hours, the presence of polyarginine leads to 2-3 times greater relative NADH (NADH_rel_ ∼ 0.3) compared to without polyarginine (NADH_rel_ ∼ 0.1) (Figure 3a, 3c). In these latter samples there was no significant difference in the reaction profiles for the two buffer conditions. However, comparison between 75 and 200 mM sodium bicarbonate with polyarginine showed that more NADH was produced at the lower concentration of sodium bicarbonate (NADH_rel_ = 0.34 ± 0.04) compared to the higher buffer conditions (NADH_rel_ = 0.25 ± 0.01). Comparison of the initial rates showed that the absence of polyarginine gave a lower initial rate (0.0073 ± 0.0008 hr^-1^ with 75 mM sodium bicarbonate and 0.0071 ± 0.0006 hr^-1^ in 200 mM sodium bicarbonate) compared to the reaction mixture with polyarginine (0.0160 ± 0.002 hr^-1^ in 75 mM sodium bicarbonate and 0.0120 ± 0.0011 hr^-1^ with 200 mM sodium bicarbonate) (Figure 3b and S17).

**Figure 3.**
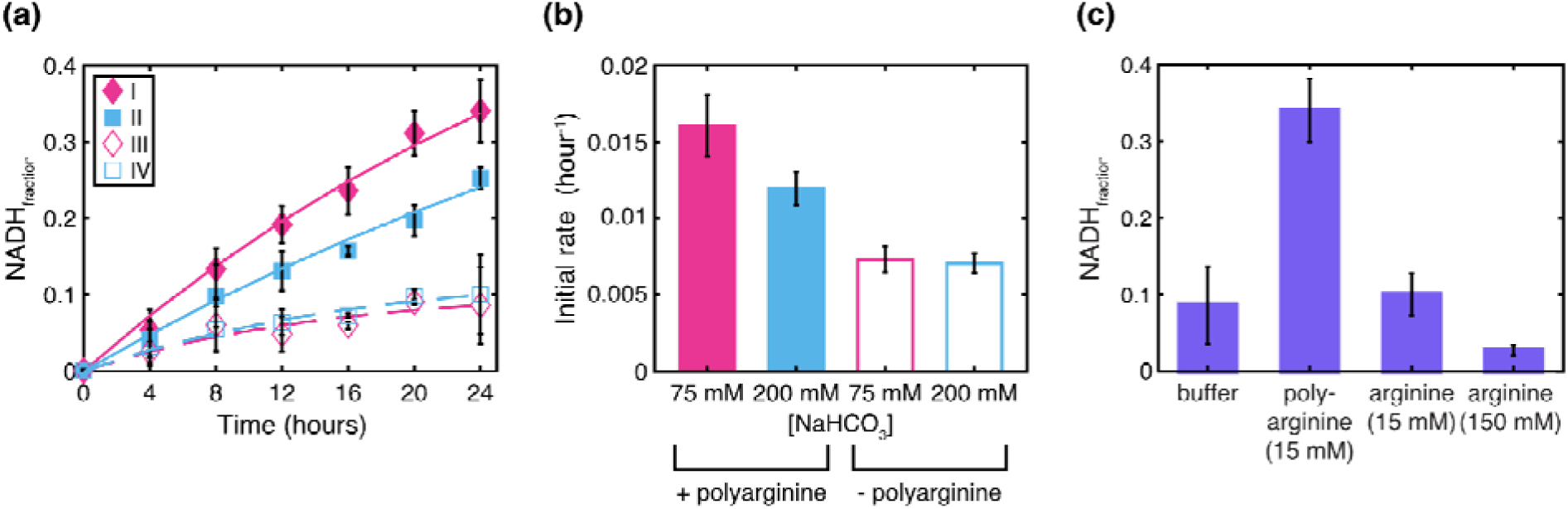
Quantification of the NADH production using commercially available biochemical assay. (a) shows the comparison of the NADH_rel_ ([NADreduced] ÷ [NADtotal]), measured at different timepoints up to 24 hours from the start of the protometabolic reaction, in the presence of poly-arginine at 75 mM sodium bicarbonate buffer (I), in the presence of poly-arginine at 200 mM sodium bicarbonate buffer (II), in the absence of poly-arginine at 75 mM sodium bicarbonate buffer (III) and in the absence of poly-arginine at 200 mM sodium bicarbonate buffer (IV). (b) shows the comparison of the initial rate at the start of the reaction calculated by fitting a straight-line, without intercept, to each of the datasets until 8 hours. (c) shows the fraction of NADH produced after 24 hours from the start of the protometabolic reaction in the presence of 75 mM sodium bicarbonate alone and with added polycations, 15 mM 50-mer polyarginine, 15 mM arginine and 150 mM arginine.

The observed differences in the fraction of NADH after 24 hours and the initial rate of NADH production could be attributed to increased charge from polyarginine rather than the phase properties. To test this, we measured the fraction of NADH at 24 hours, with monomeric arginine at 15 mM and 150 mM to simulate a charged environment in the absence of a physical droplet. In both cases we observed that the NADH fraction after 24 hours remained lower in comparison to reactions undertaken with 15 mM polyarginine that formed coacervate droplets. These results show that the increase in NADH production in the presence of polypeptide is attributed to the environment which is created by the material state. Moreover, the results show that in-situ formation of NADH modulates the material state of the dispersion from a single phase to droplets and from precipitates to droplets at different buffer conditions.

## 4. CONCLUSION

In conclusion, we used a minimal system that comprises a non-enzymatic, protometabolic electron transfer reaction with a polypeptide to test the ability for NADH to mediate different material states within an aqueous dispersion. With this simple system we demonstrate that in-situ production of NADH can drive a transition from a homogeneous solution to a dispersion with droplets and from precipitates to droplets. We propose that molecular mixtures can exhibit a variety of different phases that are stabilized by dominating interactions such as the hydrophobic effect (precipitates) and electrostatic interactions (liquid droplets) that are dependent on the chemical and physical conditions of the aqueous dispersions. We show that heterogeneous environments (droplet phases) lead to an increase in rate and amount of NADH compared to homogeneous dispersions which we attribute to the spatial rather than the chemical environment. The ability for NADH to mediate material states within aqueous protein dispersions (see Supporting Information 3, Figure S18) is important for considering how metabolism can actively tune and regulate the structural properties of the cytoplasm and how heterogeneous environments could regulate metabolism. Further, it would be pertinent to consider the consequences of an active prebiotic soup during the origin of life where carbonate-rich lakes could have provided a plausible scenario for the origins of life^31^, and concentrations of bicarbonate and salts can change upon water evaporation in wet-dry cycling scenarios^32,33^. Further, these different scenarios coupled with prebiotic chemical reactions can impact the solubility of biological relevant molecules, the chemical and material states and the outcomes of prebiotic reactions. These insights underscore the impact of early metabolic activities on the emergence of proto-cellular systems, by solubilizing precipitated polymers and leading to the formation of localized environments conducive to chemical networks, even in salt-rich environments. This active prebiotic soup potentially mirrors the properties of cytoplasm, containing various phases that can both mediate and be mediated by chemical or biochemical reactions

## ASSOCIATED CONTENT

### Supporting Information

Supplementary materials and methods; Supplementary results: NMR spectra and optical microscopy images; Initial rate fits. Addiitonal experiments: effect of NADH on chicken egg white.

## AUTHOR INFORMATION

### Present Addresses

^†^ Department of Synthetic Biology, University of Saarland, Campus Gebäude B2.2, 66123 Saarland, Germany

### Author Contributions

SM and T-YDT conceived the project; RB, AT and DR designed, conceived and undertook the experiments. RB undertook the data analysis. The manuscript was written through contributions of all authors. All authors have given approval to the final version of the manuscript.

### Notes

The authors declare no competing financial interest.

## ACKNOWLEDGMENTS

Financial support by the Horizon 2020 Marie Curie ITN (“Protomet”-Grant Agreement no. 813873 with the European Commission), MPI-CBG, ERC consolidator grant, MINSYNCELL (Grant agreement no. 101088834) gratefully acknowledged. Authors thank the ITN network PROTOMET for fruitful and lively discussions. T-YDT thanks Sangeun Lee for discussions regarding polyelectrolyte precipitation and acknowledges support from the Light microscopy facility at the MPI-CBG.

## ABBREVIATIONS

NAD^+^: nicotine adenine dinucleotide
PFK: 6-phosphofructokinase
RNA: ribonucleic acid
NMR: nuclear magnetic resonance

