## Supplementary information for "Protometabolically generated NADH mediates material properties of aqueous dispersions to coacervate microdroplets"

**Contents:**

**1. Supplementary Materials and methods**

**2. Supplementary Results**

**a) NMR**

**b) Bright field microscopy images of polyarginine phase behaviour**

**c) Determination of fraction of NADH produced**

**3. Effect of NADH on chicken egg white**

### 1. Supplementary Materials and methods

**Table S1.** Detailed list of materials used.

| **Chemical Name** | **Molecular Weight** | **Supplier** | **Product identifier** | **CAS number** |
|---|---|---|---|---|
| **2-[methoxy(polyethyleneoxy)propyltrimethoxysilane, 6-9 PEG units**<br>**(PEG-silane)** | 458.62 – 590.77 g $mol^{-1}$ | ABCR | AB111226 | 65994-07-2 |
| **β-nicotinamide adenine dinucleotide disodium salt ($NAD^{+}$)** | 685.41 g $mol^{-1}$ | Sigma Aldrich | N0632 | 20111-18-6 |
| **β-nicotinamide adenine dinucleotide, reduced disodium salt (NADH)** | 709.4 g $mol^{-1}$ | Sigma Aldrich | N8129 | 606-68-8 |
| **Deuterated Water ($D_2O$)** | 20.03 g $mol^{-1}$ | Deutero | 00506 | 7789-20-0 |
| **Guanidine thiocyanate (GuSCN)** | 118.16 g $mol^{-1}$ | PanReac AppliChem | A1107 | 593-84-0 |
| **Hydrochloric acid (HCl)** | 36.46 g $mol^{-1}$ | Merck Millipore | 1.00317 | 7647-01-0 |
| **Hellmanex III** | N / A | Hellma | 9-307-011-4-507 | N / A |
| **L-arginine monohydrochloride (arginine)** | 210.66 g $mol^{-1}$ | Sigma Aldrich | A5131 | 1119-34-2 |
| **L-lysine (lysine)** | 146.19 g $mol^{-1}$ | Sigma Aldrich | L5501 | 200-294-2 |
| **NAD/NADH quantitation kit** | N / A | Sigma Aldrich | MAK037 | N /A |

| | | | | |
|---|---|---|---|---|
| **Poly-L-arginine hydrochloride (polyarginine)** | 9.6 kDa (~ 50 mer) | Alamanda Polymers USA | PLR50 | 26982-20-7 |
| **Poly-L-lysine hydrochloride (polylysine)** | 8.2 kDa (~ 50 mer) | Alamanda Polymers USA | PLKC50 | 26124-78-7 |
| **Sodium bicarbonate ($NaHCO_3$)** | 84.01 g $mol^{-1}$ | Merck Germany | 1.06329 | 144-55-8 |
| **Sodium hydroxide (NaOH)** | 40.00 g $mol^{-1}$ | Merck Millipore USA | 1.06498 | 1310-73-2 |
| **Sodium pyruvate** | 110.04 g $mol^{-1}$ | Sigma Aldrich | P2256 | 113-24-6 |
| **Sodium trimethylsilylpropanesulfonate (DSS)** | 218.32 g $mol^{-1}$ | Tokyo Chemical Industry | T1638 | 2039-96-5 |
| **Toluene** | 92.14 g $mol^{-1}$ | Sigma Aldrich | 32249-M | 108-88-3 |

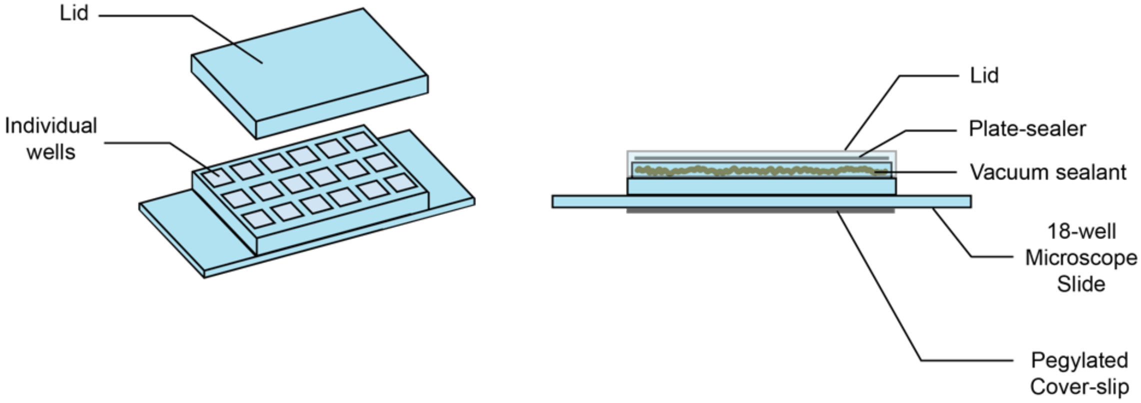


**Figure S1.** Microscope slide set-up for imaging experiments. Left: a schematic representation of the 18-well bottomless slide, consisting of a sticky underside. Right: schematic of the microscope slide that was used for time-lapse imaging.

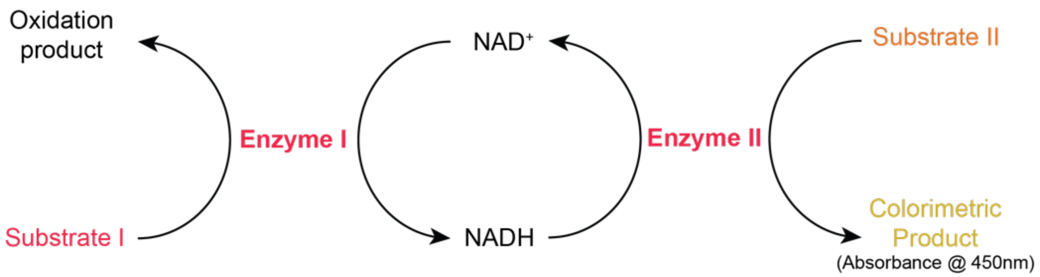


**Figure S2.** Schematic of commercially available $NAD^+$/NADH quantitation assay. The assay contains Solution 1 containing Enzymes I and II as well as the Substrate I. This enzyme mix converts $NAD^+$ to NADH. The developer solution contains Substrate II and a chromogenic substrate that converts all NADH to $NAD^+$, coupled with the conversion of the chromogenic substrate to a colorimetric product that absorbs at 450 nm.

(a)

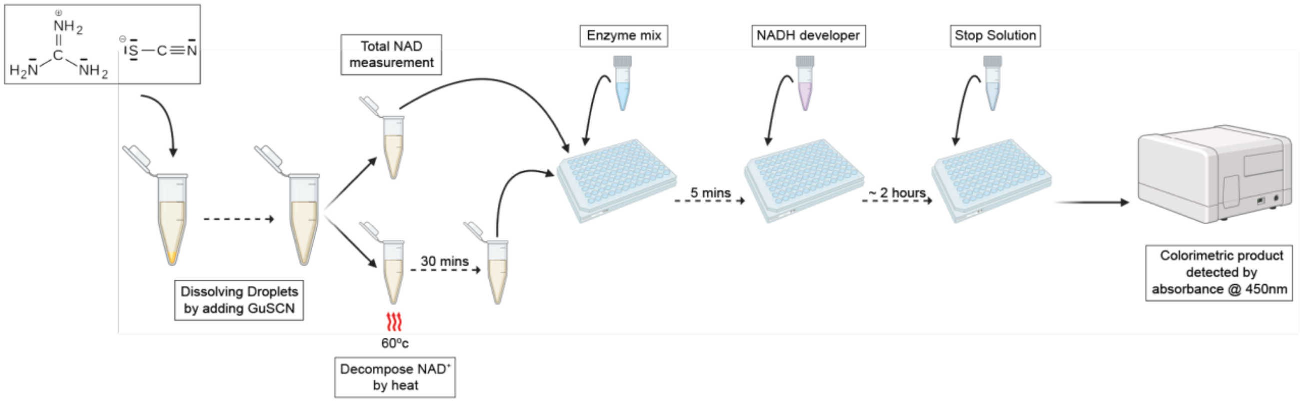


(b)

**NADH Standard Solution**

| 0 µM | 0.4 µM | 0.8 mM | 1.2 µM | 1.6 µM | 2.0 µM |
|---|---|---|---|---|---|
| 0.2284 | 0.5917 | 0.9322 | 1.3518 | 1.4303 | 1.6325 |
| 0.2263 | 0.8013 | 0.955 | 1.3826 | 1.4784 | 1.7202 |

**Sample**

| $NAD^+$ + NADH | NADH |
|---|---|
| 0.4056 | 0.2910 |
| 0.3794 | 0.3006 |

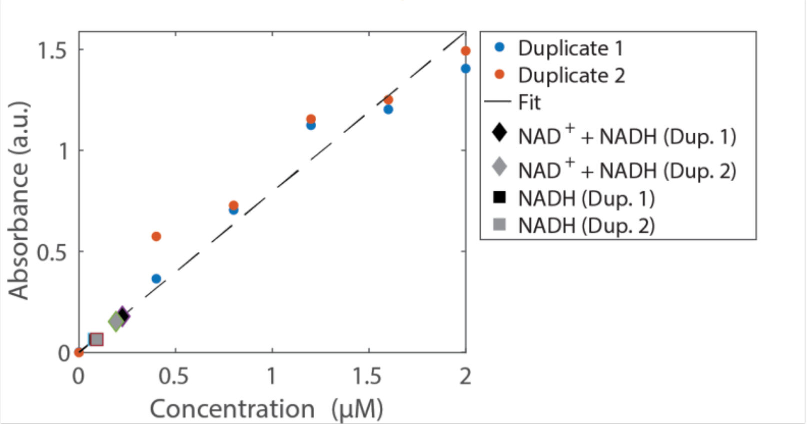


| $NAD^+$ + NADH (mM) | | NADH (mM) | | $NADH_{rel}$ | |
|---|---|---|---|---|---|
| Dup. 1 | Dup. 2 | Dup. 1 | Dup. 2 | Dup. 1 | Dup. 2 |
| 4.4883 | 3.8286 | 0.1603 | 0.1844 | 0.0357 | 0.0482 |

**Figure S3.** Schematic describing the measurement of NADH using $NAD^+$/NADH quantitation assay. (a) describes the steps involved in performing the $NAD^+$//NADH quantitation assay. First, the samples were diluted 1:1 with 4 M guanidine thiocyanate (GuSCN), to dissolve coacervate droplets. The sample was diluted in the assay buffer, supplied by the manufacturer. The sample was further diluted 1000 times and then split into two parts. One part was heated for 30 minutes at 60°C to degrade $NAD^+$, while the other part was diluted a further 10 fold and used directly in the assay. The former constitutes the sample NADH while the latter sample $NAD^+$ + NADH. To perform the assay, the enzyme mix was added to both the samples and controls containing 0 μM, 0.4 μM, 0.8 μM, 1.2 μM, 1.6 μM and 2.0 μM of NADH standard, in duplicates. This was followed by the addition of NADH developer solution after an incubation period of 5 minutes. The absorbance of the controls at 450 nm was measured in a TECAN Spark 20M wellplate reader, until the maximum value of absorbance (i.e. of 2.0 μM NADH) reached 1.0 – 2.0. A stop solution, also supplied by the manufacturer, was then added to arrest the enzymatic activity and prevent the degradation of the colorimetric product. After the addition of the stop solution the samples and the controls were then measured for absorbance at 450 nm. (b) shows the detailed steps of the data analysis for a single sample, as an illustrative example. The absorbance values of the controls were plotted against the corresponding NADH concentration. This is then linearly fitted, using least squares' method, to the straight line Y = mX, where Y is the absorbance and X is the corresponding concentrations, to obtain the slope, m. Using the slope, the concentration of NADH and $NAD^+$ + NADH was obtained for each duplicate as concentration = (absorbance / slope) × dilution factor. The amount of NADH was reported as $NADH_{rel}$, such that for each duplicate $NADH_{rel}$ = NADH (mM) ÷ (mean of 2 duplicates of $NAD^+$ (mM) + NADH (mM)).

### 2. Supplementary Results

#### a) NMR

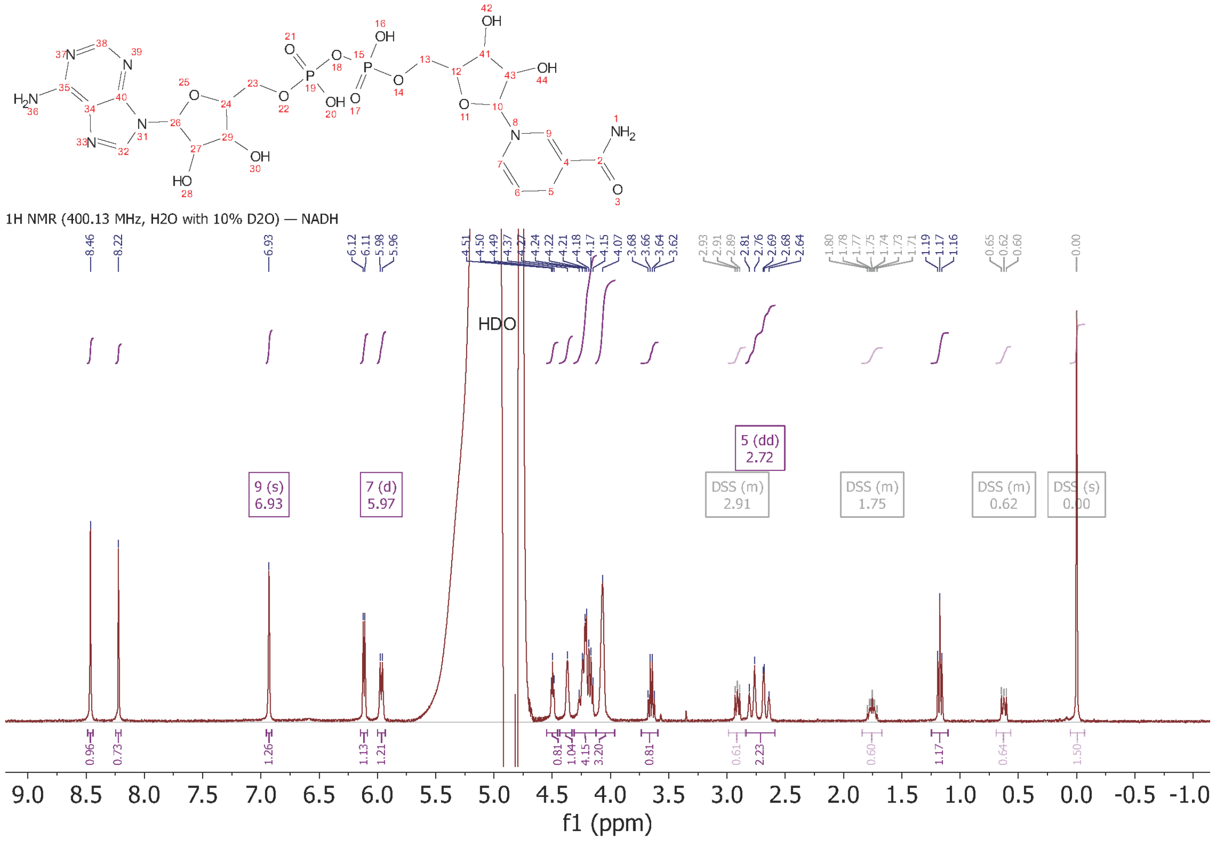


**Figure S4.** $^{1}H$ NMR spectra of NADH (400 MHz, $H_2O$ with 10% $D_2O$) with DSS internal standard. The assignable hydrogens have been marked on the spectra. For reference the structure is provided; δ 2.72 (dd, $J$ = 14.3, 7.9 Hz, 2H), 5.97 (d, $J$ = 8.2 Hz, 1H), 6.93 (s, 1H).

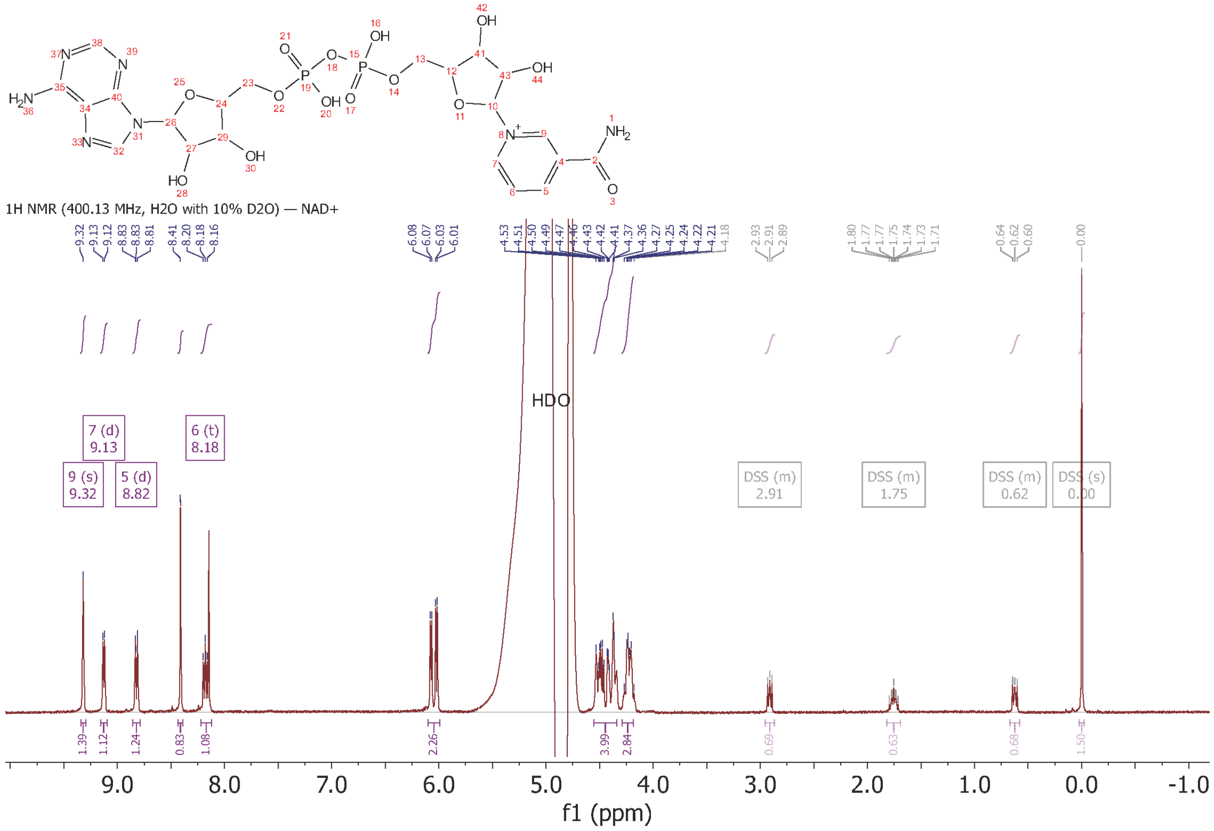


**Figure S5.** $^{1}$H NMR spectra of NAD$^{+}$ (400 MHz, $H_2O$ with 10% $D_2O$) with DSS internal standard. The assignable hydrogens have been marked on the spectra. For reference the structure is provided; δ 8.18 (t, $J$ = 7.2, 7.2 Hz, 1H), 8.82 (d, $J$ = 8.1 Hz, 1H), 9.13 (d, $J$ = 6.3 Hz, 1H), 9.32 (s, 1H).

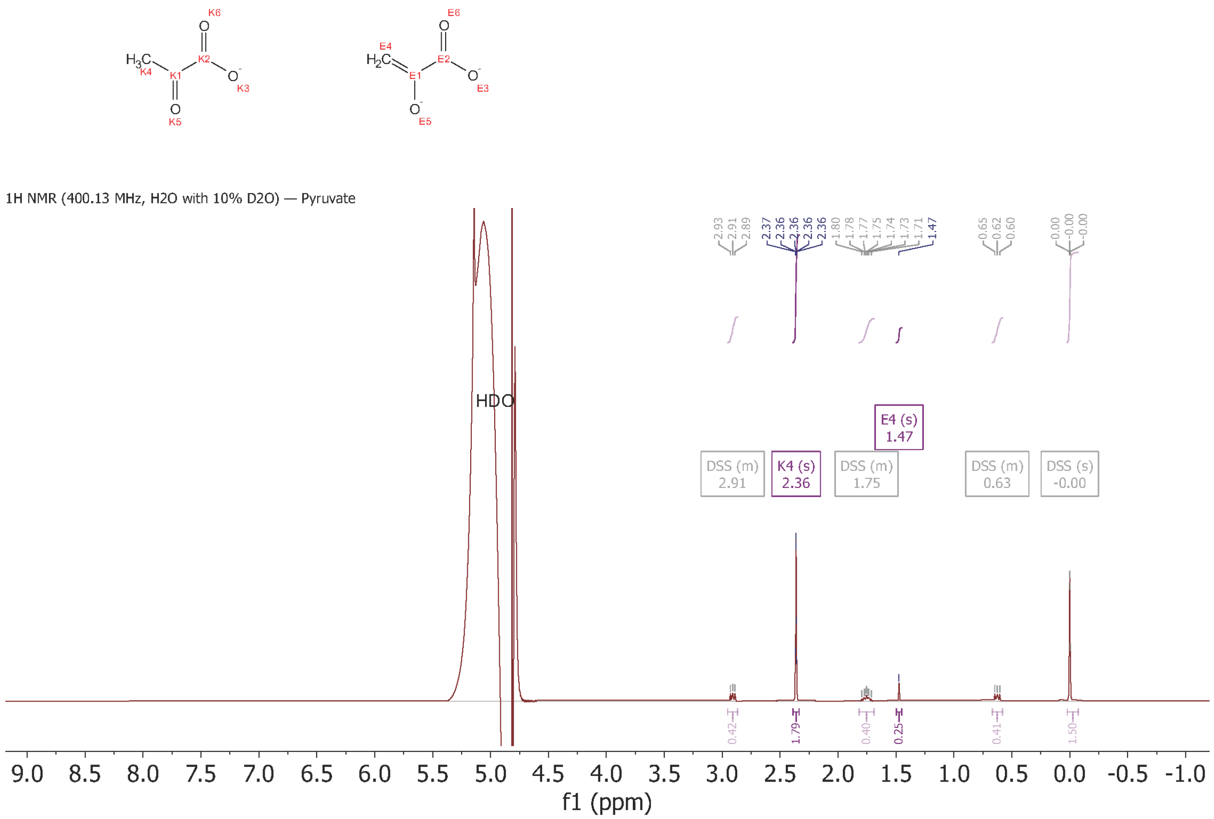


**Figure S6.** $^1$H NMR spectra of pyruvate (400 MHz, $H_2O$ with 10% $D_2O$) with DSS internal standard. The assignable hydrogens have been marked on the spectra. The structures is provided for reference; δ 1.47 & 2.36 (s, 3H).

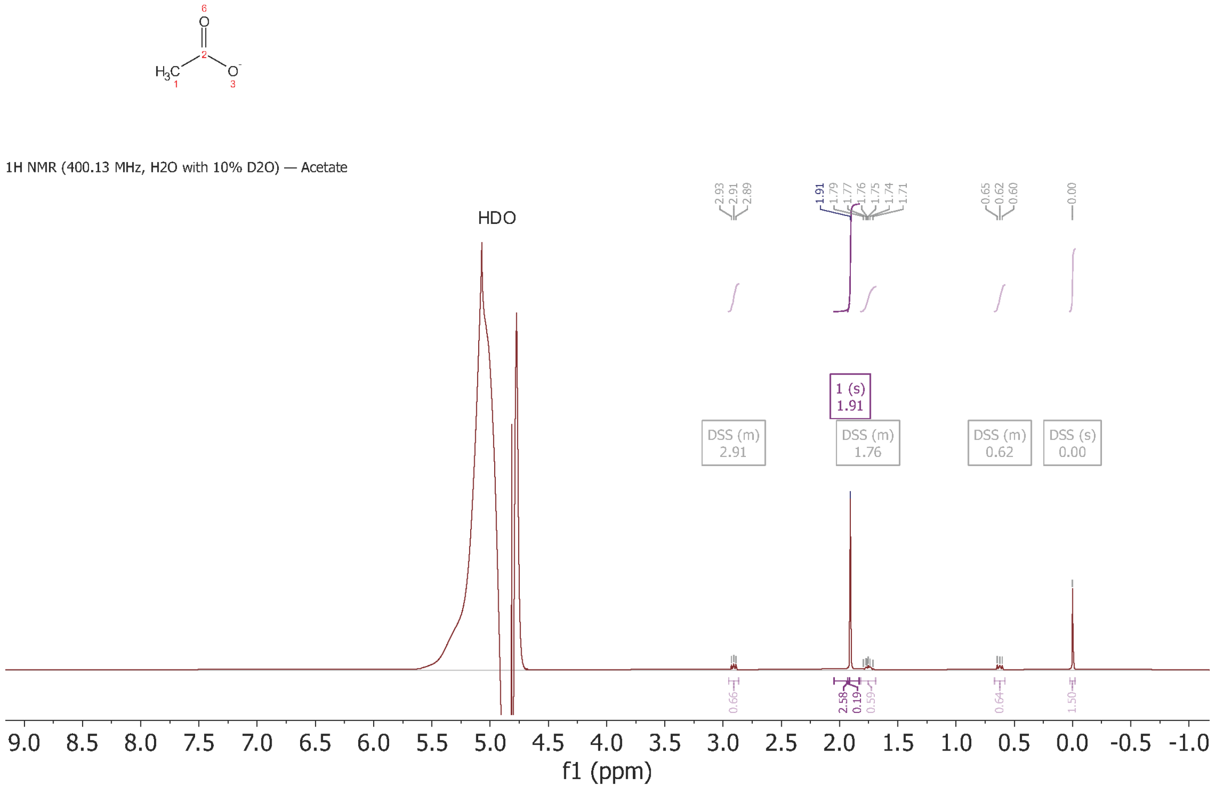


**Figure S7.** $^{1}H$ NMR spectra of Acetate (400 MHz, $H_2O$ with 10% $D_2O$) with DSS internal standard. The assignable hydrogens have been marked on the spectra. The structure is provided for reference; δ 1.91 (s, 3H).

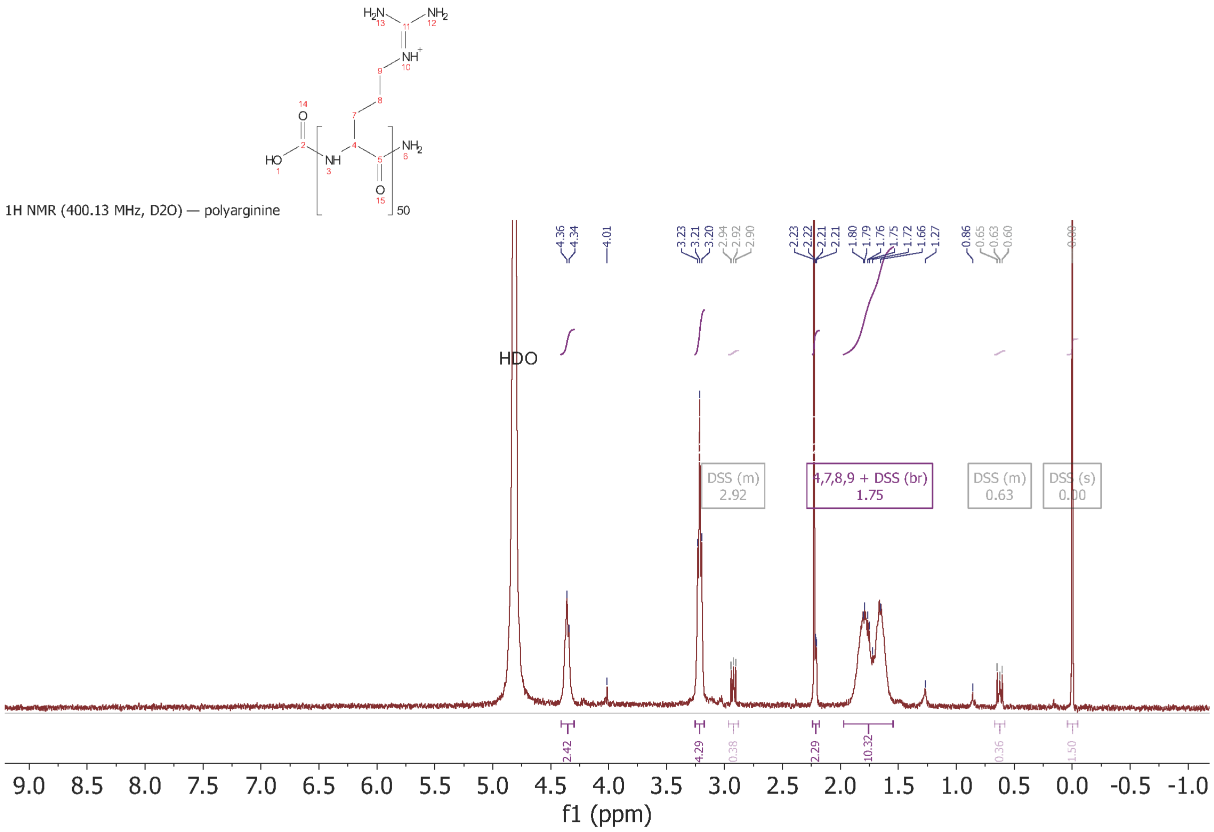


**Figure S8.** $^{1}$H NMR spectra of polyarginine (400 MHz, $D_2O$) with DSS internal standard. The assignable hydrogens have been marked on the spectra. For reference the structure is provided; δ 1.97 – 1.55 (br).

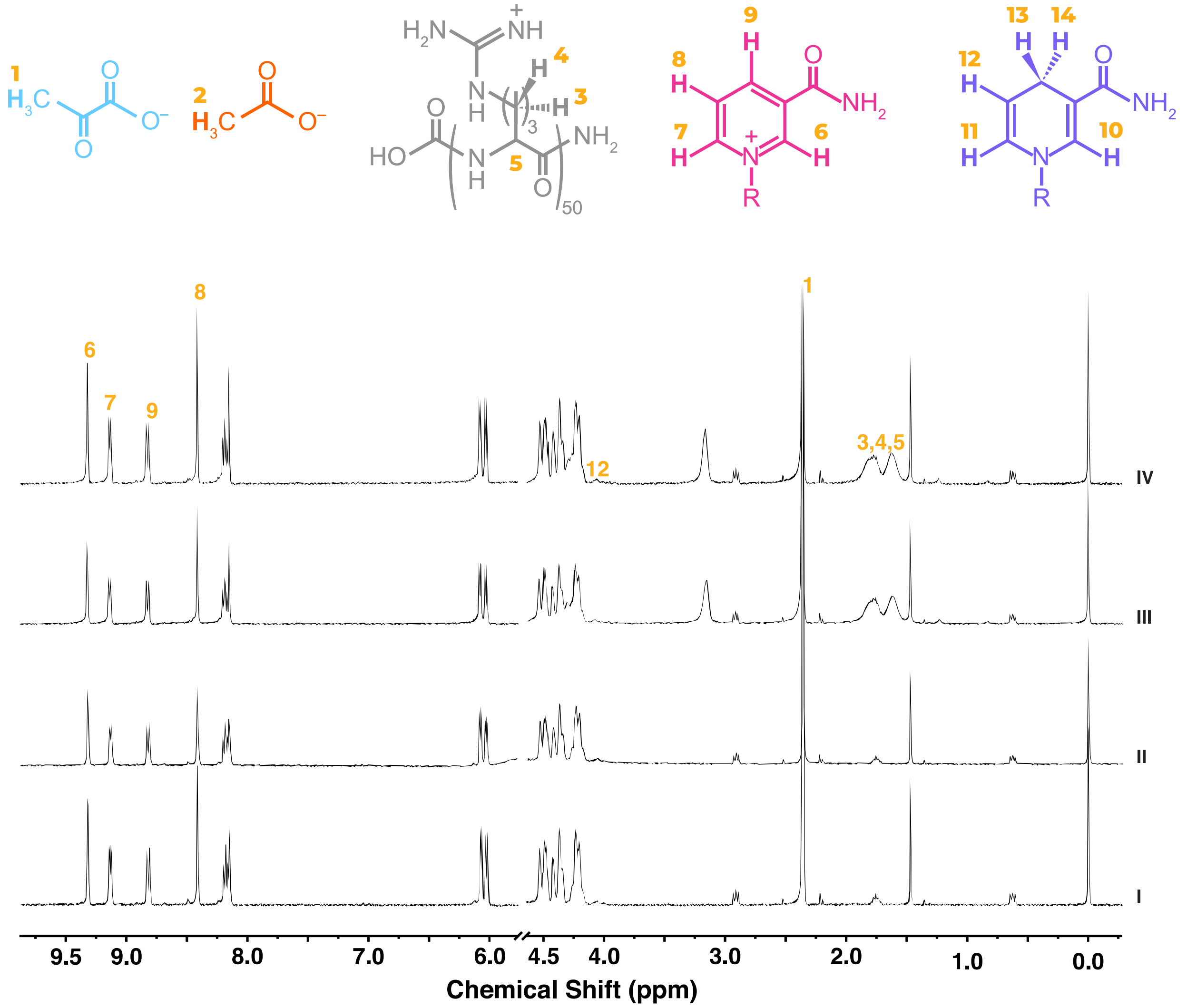


**Figure S9.** $^{1}H$ NMR spectra of reaction mixtures at the start of the reaction. I, II, III and IV are reaction mixtures without polyarginine in 75 mM $NaHCO_3$ buffer, without polyarginine in 200 mM $NaHCO_3$ buffer, with 15 mM polyarginine in 75 mM $NaHCO_3$ buffer and with 15 mM polyarginine in 200 mM $NaHCO_3$ buffer.

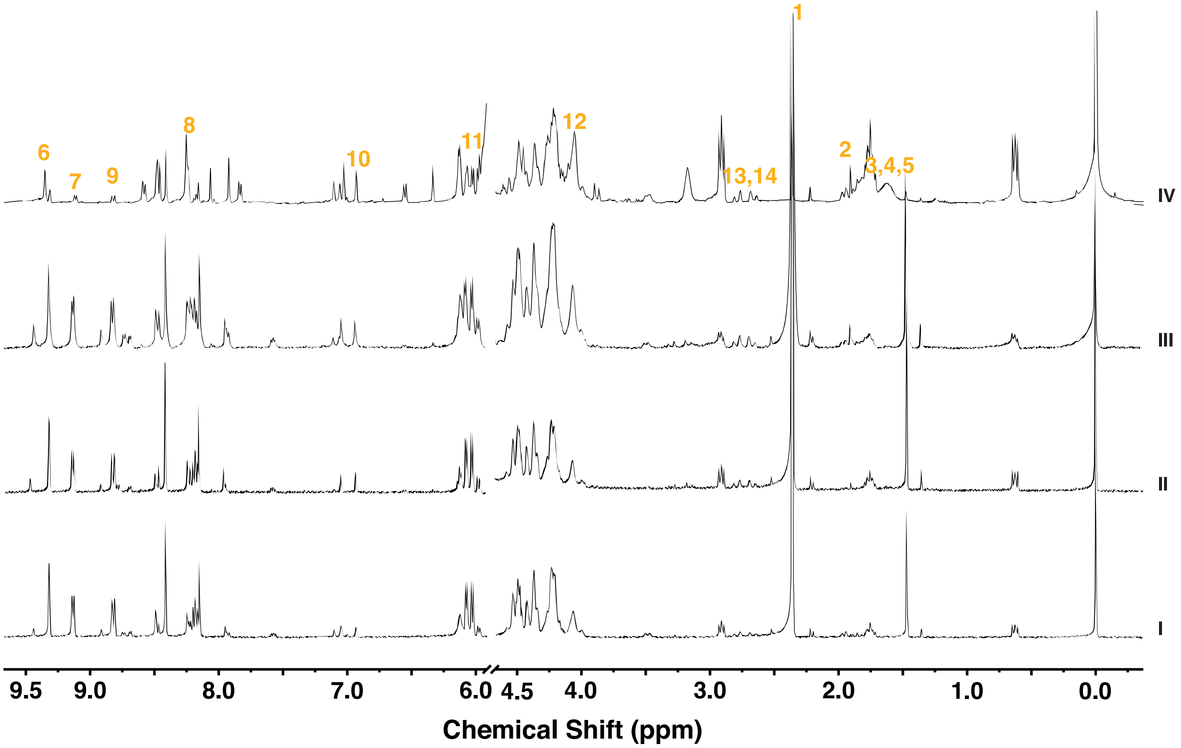


**Figure S10.** $^1$H NMR spectra of reaction mixtures prepared in 75 mM $NaHCO_3$ buffer after 24 hours of the reaction. I and II are reaction mixtures without polyarginine and with 15 mM polyarginine. Reaction mixtures with 15 mM polyarginine was centrifuged to separate the dilute phase and the coacervate phase. III and IV the $^1$H-NMR spectra of the dilute and the coacervate phase redissolved in 75 mM $NaHCO_3$ buffer.

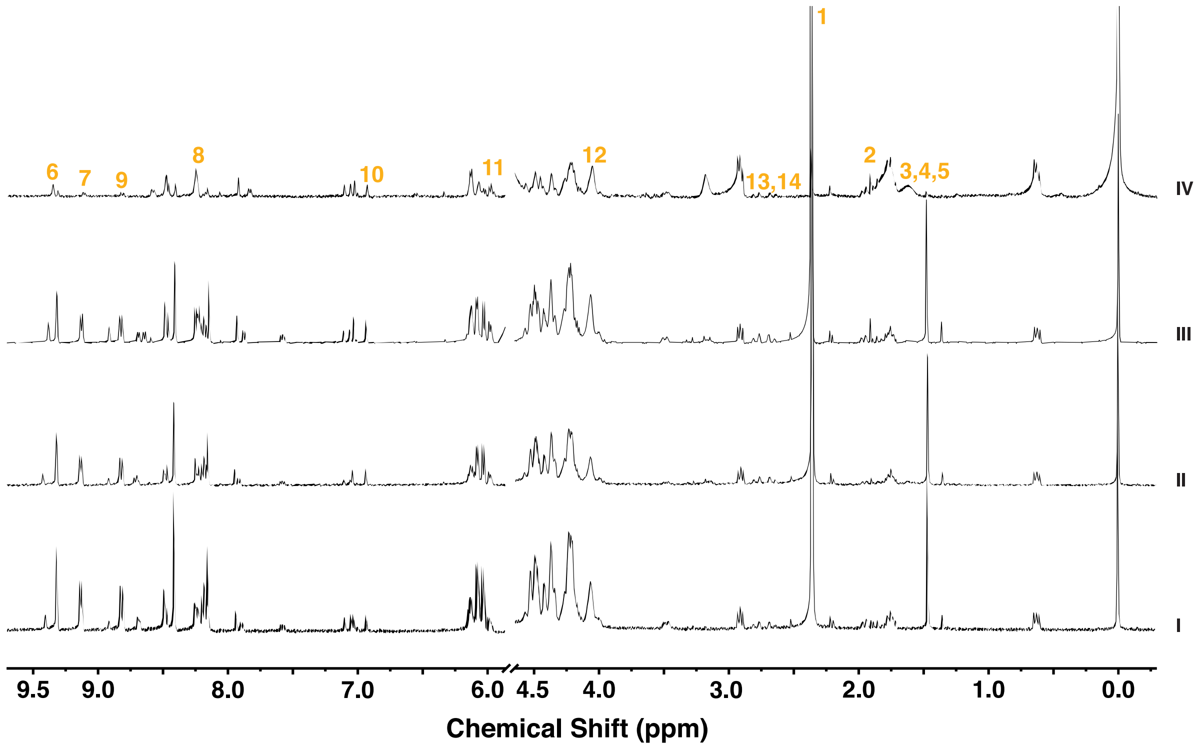


**Figure S11.** $^1H$ NMR spectra of reaction mixtures prepared in 200 mM $NaHCO_3$ buffer after 24 hours of the reaction. I and II are reaction mixtures without polyarginine and with 15 mM polyarginine. Reaction mixtures with 15 mM polyarginine was centrifuged to separate the dilute phase and the coacervate phase. III and IV the $^1H$-NMR spectra of the dilute and the coacervate phase redissolved in 200 mM $NaHCO_3$ buffer.

b) **Bright field microscopy images of polyarginine phase behaviour**

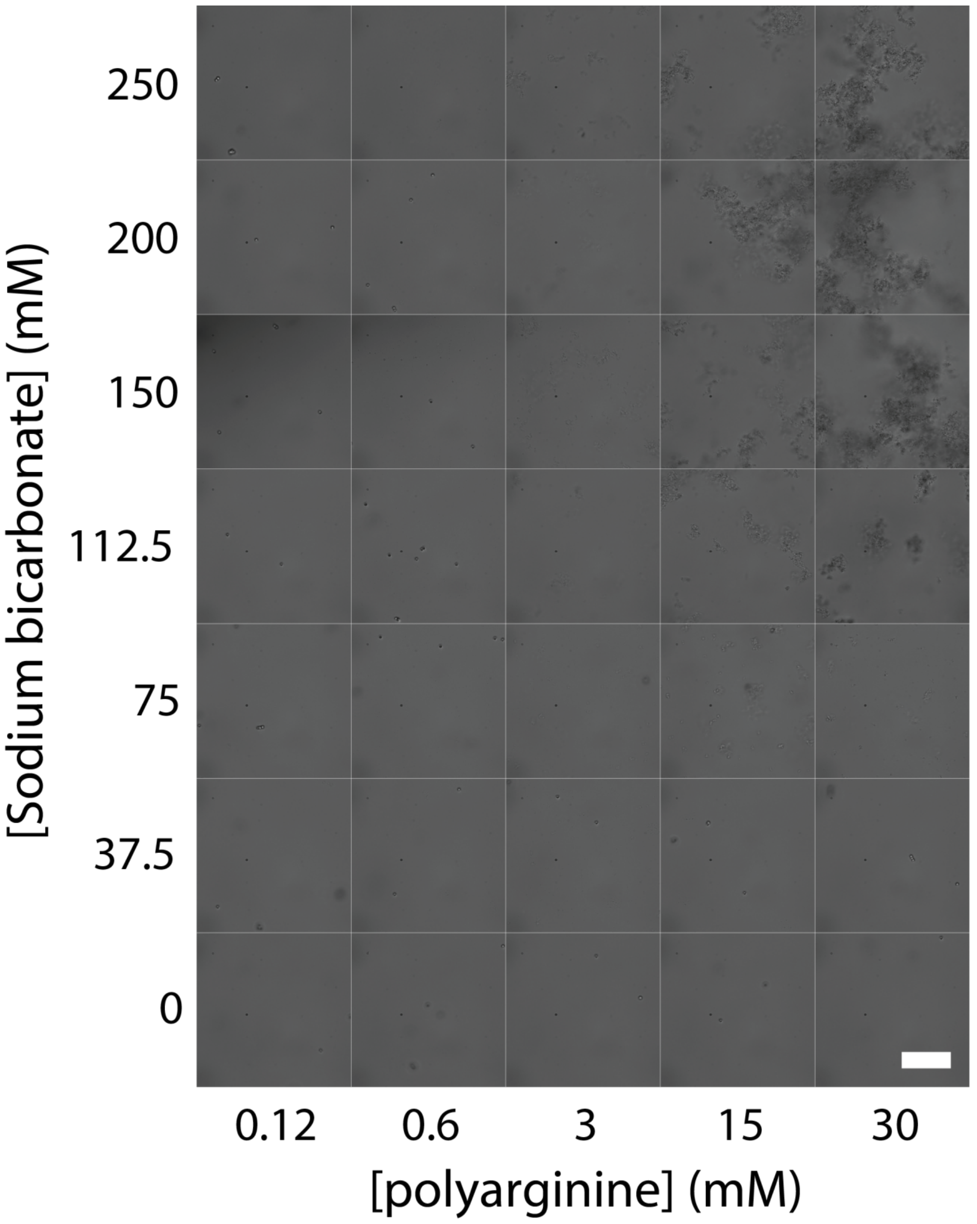


**Figure S12.** Bright field microscopy images mixtures of sodium bicarbonate and 50-mer polyarginine showing their phase behaviour, comprising of either dissolved or precipitated state. To aid visualisation, carboxylate microspheres were added to samples that did not contain precipitates. Scale bar = 20 µm.

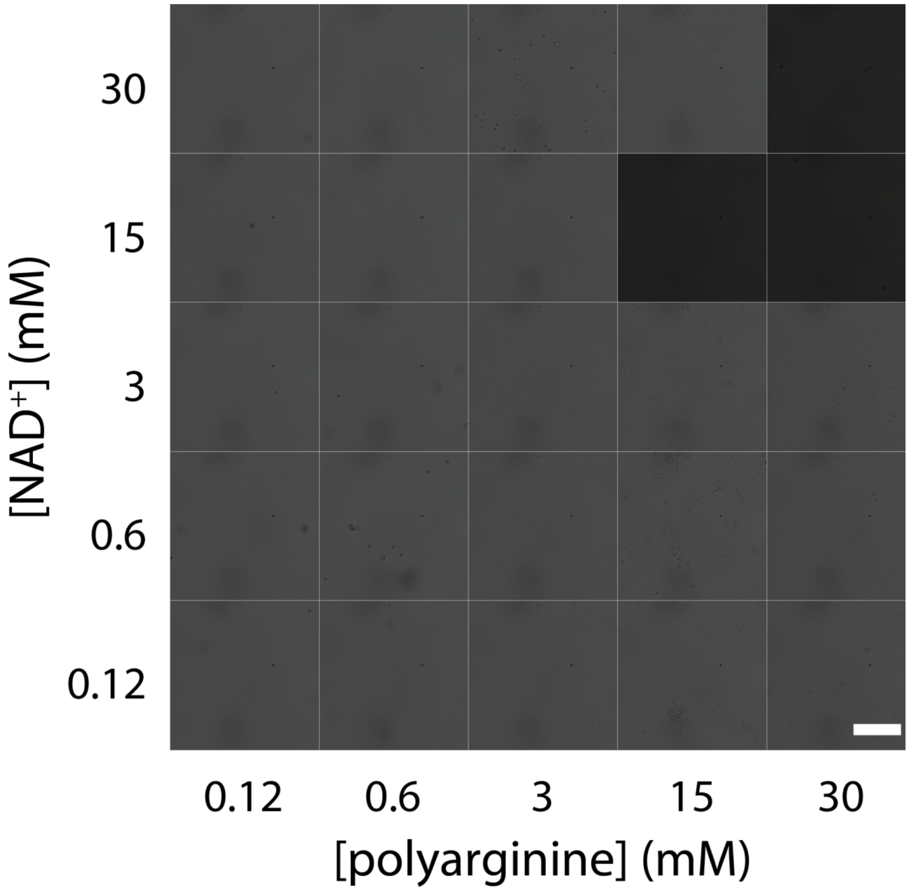


**Figure S13.** Bright field microscopy images mixtures of $NAD^+$ and 50-mer polyarginine, in 75 mM sodium bicarbonate, showing their phase behaviour, comprising of either dissolved, coacervated or precipitated state. To aid visualisation, carboxylate microspheres were added to samples that did not contain precipitates or coacervates. Scale bar = 20 µm.

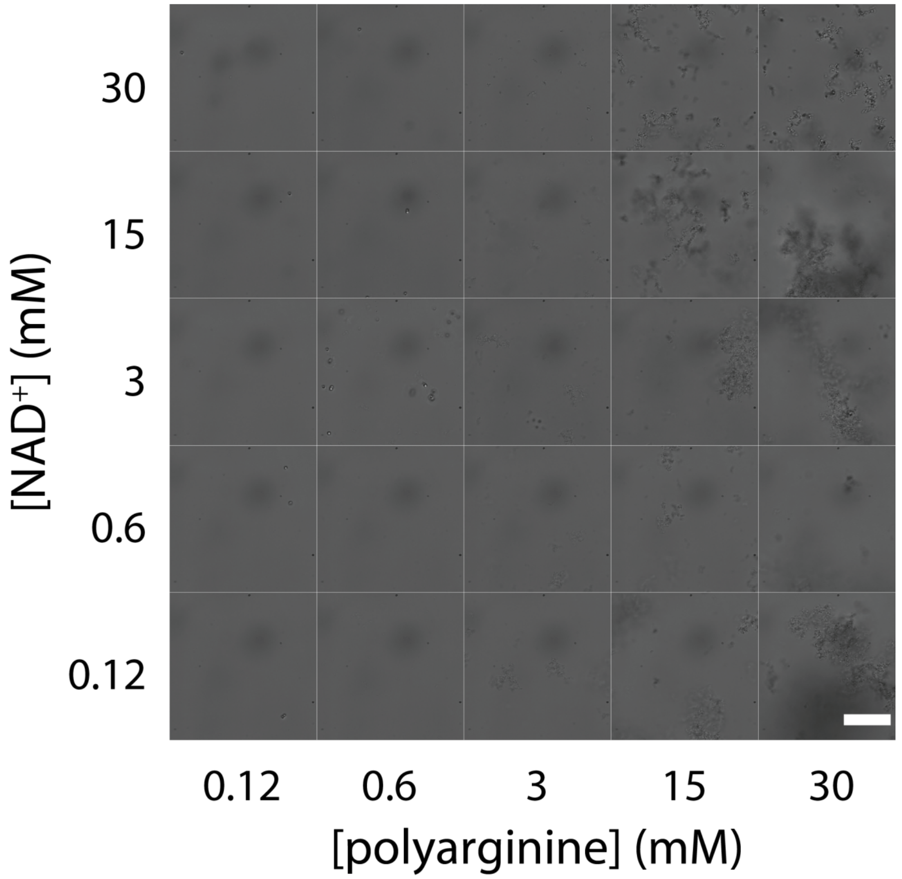


**Figure S14.** Bright field microscopy images mixtures of $NAD^+$ and 50-mer polyarginine, in 200 mM sodium bicarbonate, showing their phase behaviour, comprising of either dissolved or precipitated state. To aid visualisation, carboxylate microspheres were added to samples that did not contain precipitates. Scale bar = 20 μm.

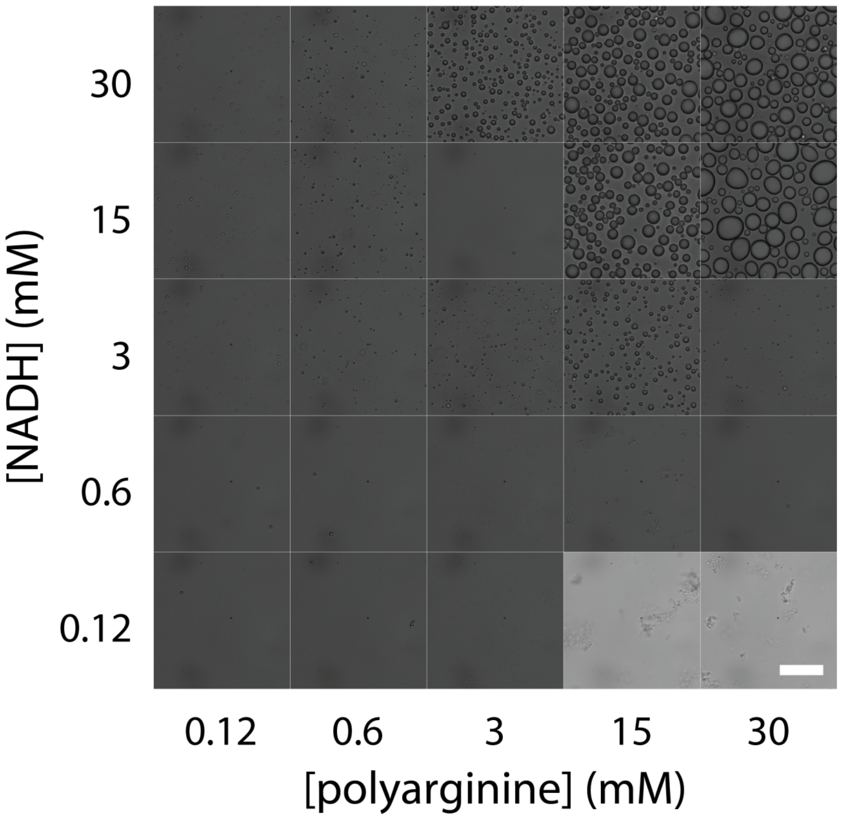


**Figure S15.** Bright field microscopy images mixtures of NADH and 50-mer polyarginine, in 75 mM sodium bicarbonate, show regions of homogeneity, droplets and precipitates. To aid visualisation, carboxylate microspheres were added to samples that did not contain precipitates or coacervates. Scale bar = 20 µm.

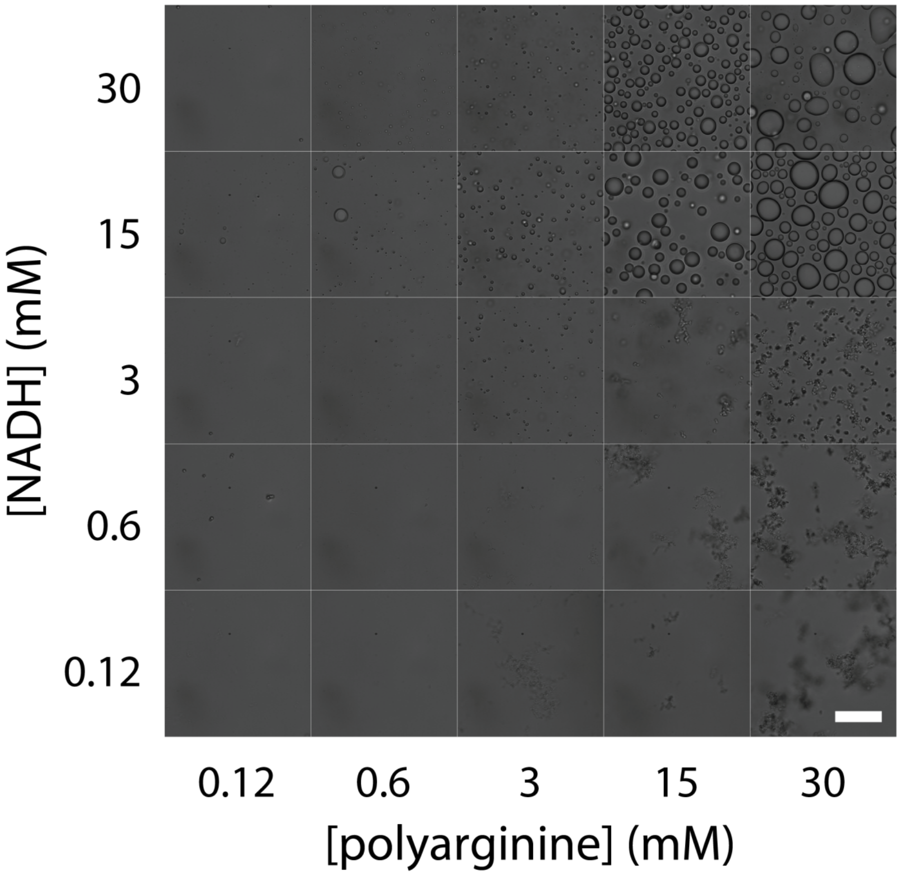


**Figure S16.** Bright field microscopy images mixtures of NADH and 50-mer polyarginine, in 200 mM sodium bicarbonate, showing their phase behaviour, comprising of either dissolved, coacervated or precipitated state. To aid visualisation, carboxylate microspheres were added to samples that did not contain precipitates or coacervates. Scale bar = 20 µm.

c) **Determination of fraction of NADH produced**

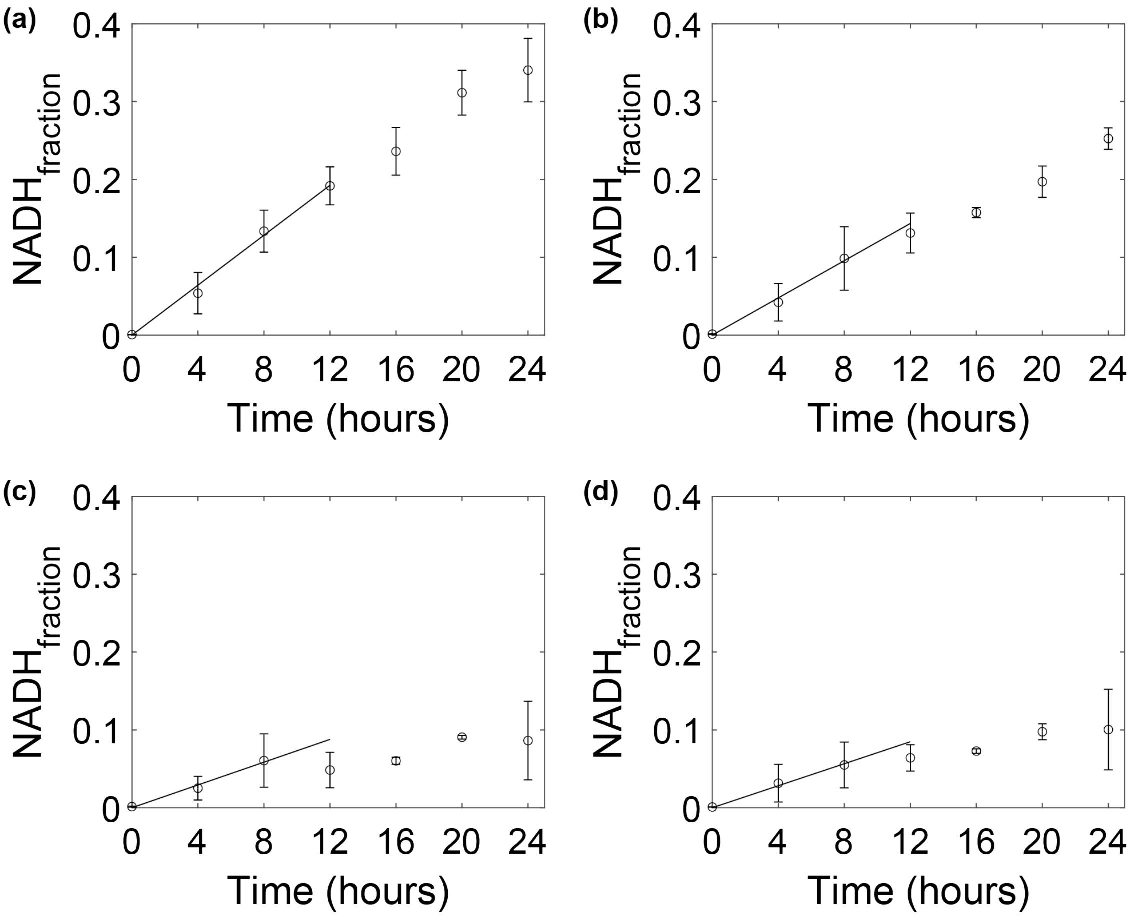


**Figure S17.** Linear least squares fit of NADH fraction to obtain the initial rate. Panels (a), (b), (c), and (d) show the $NAD^+$ reduction in the presence of 50-mer polyarginine in 75 mM sodium bicarbonate, 50-mer polyarginine in 200 mM sodium bicarbonate, only 75 mM sodium bicarbonate, and only 200 mM sodium bicarbonate, respectively. The slope of each fitted line indicates the initial reaction rate: 0.0160 ± 0.002, 0.0120 ± 0.0011, 0.0073 ± 0.0008, and 0.0071 ± 0.0006 $hr^{-1}$, respectively. The slope of fit has been extended from 8 to 12 hours for visualization. The fit comes from fitting to the first 8 hours. Error in the data points are the standard deviation from at least 3 repeats, the errors are calculated based on a 95% confidence interval.

### 3. **Effect of NADH on chicken egg white albumin**

To test the effect of $NAD^+$ and NADH on chicken egg white medium we used a protocol as previously described[1]. Medium-sized, free-range chicken eggs were purchased from a supermarket and utilized before the stated expiration date. The egg white was separated from the yolk by careful shell-to-shell transfer. The chicken egg white was diluted 2-fold with 50 mM Tris/HCl buffer (50 mM, pH 7.4) and centrifuged at room temperature for 10 minutes at 5,000 × g. The supernatant was further diluted 5-fold with Tris/HCl buffer to a final concentration of 40 mM Tris-HCl. 40 µL of $NAD^+$, NADH prepared in TRIS-HCl buffer (50 mM, pH 7.4) were added to the 160 µL of supernatant to achieve a final concentration of 10 mM of NaCl, $NAD^+$ or NADH) and 42 mM Tris-HCl in a 0.2 mL thin-walled PCR tubes. The samples were incubated at 60°C in an Applied Biosystems™ MiniAmp™ Plus Thermocycler.  As a control experiment, 40 µL of Tris-HCl was added to the diluted chicken egg white albumin. Images were captured on a black background with a smartphone camera.

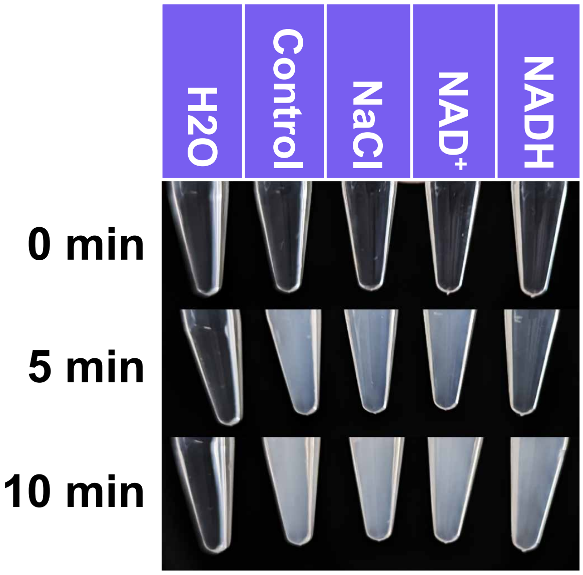


**Figure S18.** Images showing the increase in albumin turbidity with increasing time. Results show that NADH delays the onset of increased turbidity at 5 mins. 10 mM of NaCl, $NAD^+$, NADH or water were added to diluted chicken egg white to a final concentration of 10 mM for NaCl, $NAD^+$ and NADH respectively.
